# Mass spectra alignment using virtual lock-masses

**DOI:** 10.1101/292425

**Authors:** Francis Brochu, Pier-Luc Plante, Alexandre Drouin, Dominic Gagnon, Dave Richard, Francine Durocher, Caroline Diorio, Mario Marchand, Jacques Corbeil, François Laviolette

## Abstract

Mass spectrometry is a valued method to evaluate the metabolomics content of a biological sample. The recent advent of rapid ionization technologies such as Laser Diode Thermal Desorption (LDTD) and Direct Analysis in Real Time (DART) has rendered high-throughput mass spectrometry possible. It is used for large-scale comparative analysis of populations of samples. In practice, many factors resulting from the environment, the protocol, and even the instrument itself, can lead to minor discrepancies between spectra, rendering automated comparative analysis difficult. In this work, a sequence/pipeline of algorithms to correct variations between spectra is proposed. The algorithms correct multiple spectra by identifying peaks that are common to all and, from those, computes a spectrum-specific correction. We show that these algorithms increase comparability within large datasets of spectra, facilitating comparative analysis, such as machine learning.

## Introduction

Mass spectrometry (MS) is a widely used technique for acquiring data on the metabolome or the proteome of individuals^12^. Proteomics applications can consist, among others, of typing of microbial organisms^3^, imaging MS^4^, quantitative comparisons^5^, and peptide sequencing^67^. For metabolomics applications, the two main approaches fall into the categories of targeted and untargeted studies. In comparison with targeted studies, untargeted studies acquire data using a shotgun approach. Therefore, this type of study is a good option for novel biomarker discovery and hypothesis generation^89^.

Through recent years, novel ionization technologies have emerged, facilitating the high-throughput acquisition of mass spectra^10^. Technologies such as Laser Diode Thermal Desorption (LDTD) or Direct Analysis in Real Time (DART), allow for the rapid acquisition of large datasets. These methods often preclude or bypass the time separation process used in Liquid Chromatography (LC) or Gas Chromatography (GC)^11^. Thus, without any time separation, a single mass spectrum will often be represented as lists of peaks, composed of the mass-to-charge ratio of the ion (*m*/*z* value) and its intensity. Although, in this case, a single mass spectrum will be more complex and composed of more peaks and compounds than a spectrum with a time-separation method.

With the rise of larger datasets, multiple problems of comparability between spectra have emerged. Datasets are acquired in multiple batches over numerous days, on different instruments in multiple locations, with recalibrations of the instruments occurring between batches^12^. These factors induce variations in the spectra that hinders their comparison.

In the past, three algorithms have been proposed to address this problem, mainly affecting Time-of-Flight mass spectrometers. These include the work of Tibshirani *et al.* (2004)^13^, Jeffries (2005)^14^, and Tracy *et al.* (2008)^15^. Tibshirani’s algorithm relies on a clustering algorithm to align peaks that are present in multiple spectra and picks them for further statistical analyses. However, unlike the algorithms proposed in this article, it does not address the problem of inter-batch variations. Jeffries’ algorithm is more appropriate for this problem. This method uses cubic splines to recalibrate spectra, based on the shifts between observed peaks and known reference masses. A similar algorithm has been proposed by Barry *et al.* (2013) for Fourier-Transform Mass Spectrometry^16^. This approach uses ambient ions in order to correct the spectra using known reference masses. One limitation of these algorithms is that they require known reference masses. The algorithm presented in this work alleviates this constraint, by automatically detecting such reference points. Another algorithm of interest for MALDI-ToF spectra has been proposed by Tracy *et al.*^15^. In this case, commonly occurring peaks within the dataset are used to correct the spectra and determine the binning distance used. However, this method computes a single constant correction factor for the entire spectrum, while the method proposed in this work computes correction factors that vary across the *m*/*z* axis of the spectra in order to obtain a more accurate correction.

The algorithms proposed in this article aim to render spectra more comparable prior to peak selection and statistical analyses. We draw inspiration from the internal lock mass approach and exploit the fact that spectra of samples of the same nature (i.e., blood plasma samples, urine samples, etc.) are very likely to share common peaks (i.e., compounds that are present in each sample). The internal lock mass approach consists of introducing a standard compound along with the sample into the ion source^17^. T̊his known compound can then be used to correct instrumental drift in *m*/*z* values, potentially in real-time. For example, human blood plasma contains compounds, such as glucose and amino acids^18^. Similarly, urine contains urea, creatinine, citric acid, and many more^19^. Hence, we propose to correct the spectra based on the position of peaks that are detected to be consistently present in samples of the same nature. We call these peaks “virtual lock masses” (VLM) and propose an algorithm to detect them automatically. This idea is similar to the one proposed by Barry *et al.*^16^, but the peaks are not limited to ambient ions. In this work, we show that our algorithm allows the detection of tens to hundreds of peaks that can be used as reference points to re-align the spectra. These points will serve to reduce inter-batch and intra-batch variations in the spectra, but will not correct the spectra to the true *m*/*z* values of the ions. However, our approach is fully compatible with the classical lock mass approach, which can be used complementarily. Moreover, we show that a slight modification to the VLM detection algorithm can produce an alignment algorithm that can be used to further correct the spectra.

Hence, our key contributions are: an algorithm that automatically detects reference points in mass spectra, an algorithm that corrects the spectra based on these points, and an alignment algorithm to align large sets of spectra. The next section describes the details of the algorithms and their implementation and we then present results supporting the accuracy of our reference point detection algorithm. Moreover, we show that the proposed algorithms yield classifiers with increased accuracy in the context of machine learning analysis performed on ToF mass spectra. Finally, we discuss these results and their implications.

## Methods

In this section, we present the mathematical basis of the proposed methodology. First, the problem of virtual lock-mass identification is addressed. A formal definition of VLM peaks is introduced, along with an highly efficient algorithm capable of identifying such peaks in a set of mass spectra. Second, a methodology for correcting mass spectra based on a set of identified virtual lock masses is described. Third, an algorithm for mass spectra alignment based on the previous algorithm is proposed. Finally, the datasets used and the experimental methodologies are presented. Note that the algorithms are designed to be applied to partially pre-processed spectra, specifically processed to centroid format.

### Definitions

Let us first recall that a *set* is an un-ordered collection of elements whereas a *sequence* is an ordered collection of elements. Hence, in a sequence we have a first element, a second element, and so on. If *A* is a sequence or a set, |*A*| denotes the number of elements in *A*.

Let 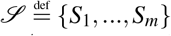 be a set of mass spectra. Each spectrum *S*_*i*_ is a sequence of peaks, where each peak is a pair (*μ, ι*) with an *m*/*z* value *μ* and a peak intensity *ι*. Let a *window* of size 2*w* centered on the peak (*μ*, *ι*) be an interval that starts at *μ* · (1 − *w*) and ends at *μ* · (1 + *w*). Notice that the size of the window *w* is relative to *μ*. The reason for using window sizes in relative units is that the mass measurement uncertainty of ToF mass spectrometers increases linearly with the *m*/*z* value of a peak.

Given a set 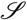 of mass spectra and a window size parameter *w*, a *virtual lock mass* (VLM) with respect to (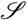, *w*) is a point *v* on the *m*/*z* axis such that there exists a set 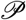 of peaks from 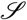 that satisfies the following properties

1. 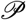 contains exactly one peak from each spectrum in 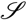.
2. The average of the *m*/*z* values of the peaks in 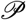 is equal to *v*.
3. Every peak in 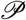 has a *m*/*z* value located in the interval [*v*(1 − *w*), *v*(1 + *w*)].
4. No other peak in 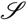 has an *m*/*z* value that belongs to [*v*(1 − *w*), *v*(1 + *w*)].
5. Every peak in 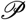 has an intensity in the interval [*t*_*a*_,*t*_*b*_].

If and only if all these criteria are satisfied, we say that 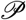 is the set of peaks associated with the VLM *v*.

Note that we impose a lower intensity threshold *t*_*a*_, since peaks with a lower intensity will tend to have a lower mass accuracy, and can even be confused with noise. In addition, there is also accuracy issues when the intensity of a peak is higher than the machine specifications. Consequently, we also impose an upper intensity threshold *t*_*b*_. Hence, in principle, a VLM is defined only with respect to (w.r.t.) (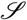, *w*, [*t*_*a*_,*t*_*b*_]). However, we will drop the reference to *t*_*a*_ and *t*_*b*_ to simplify the notation.

A crucial aspect of the definition of a VLM is the fact that it holds only w.r.t. a given window size *w*. Indeed, consider Fig (1) which represents peaks coming from three different spectra. We can observe that a first window size *w*_1_ will correctly detect four VLM points. If the window size is too large however, we observe the case of *w*_2_: peaks that are further apart can be erroneously grouped into a VLM group. Moreover, *w*_2_ can detect the first grouping of peaks within the figure as a VLM, and then the shown grouping as a second one. Thus, the same peaks would be part of two distinct VLM points. This would create ambiguity in the correction and is nonsensical. The last possible case is that of a window size that is too small. In this situation, the window would be unable to detect groups of peaks coming from each spectra of 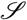.

**Figure 1.**
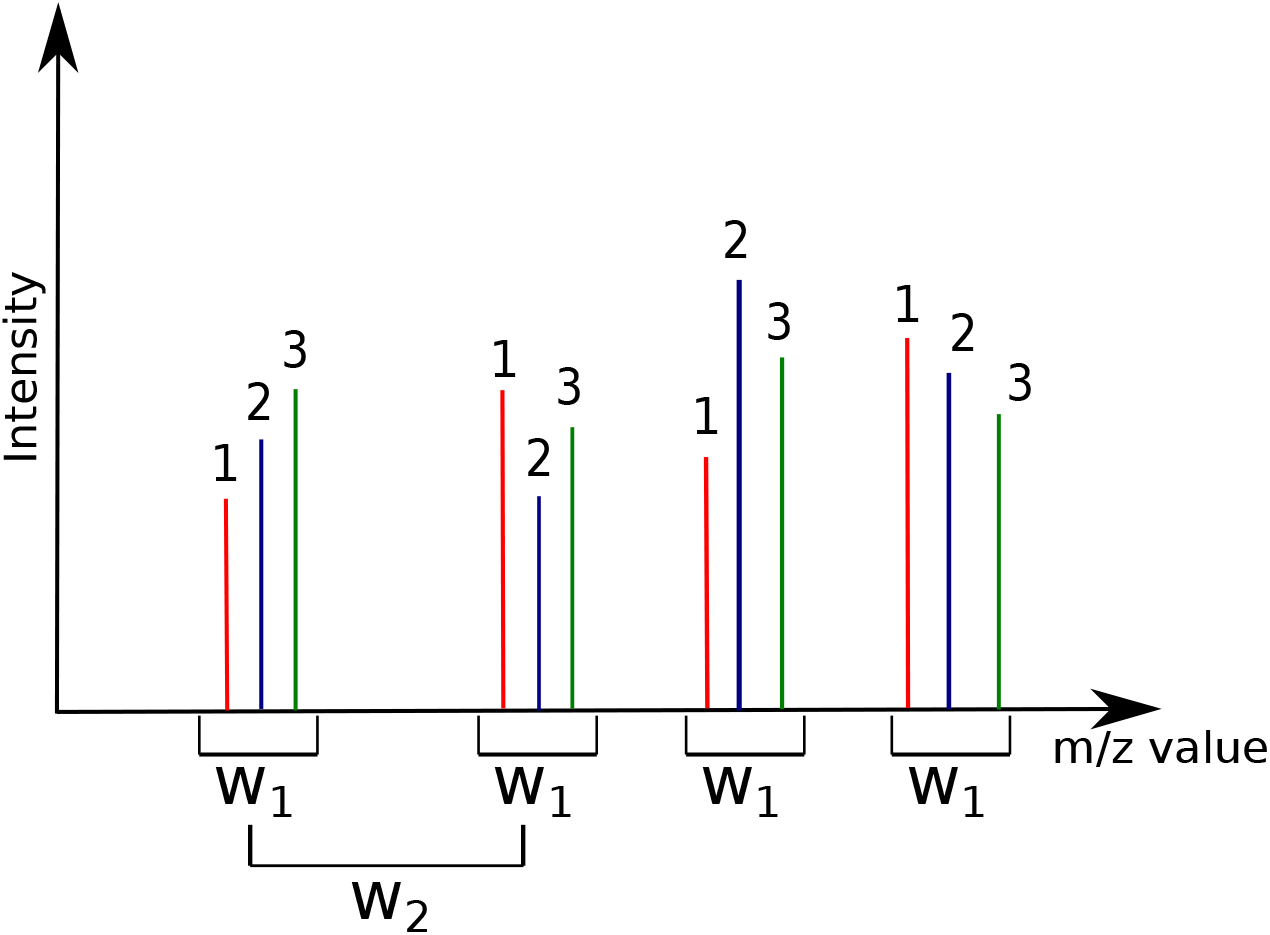
Definition of window size for the detection of VLM peaks. The peaks identified as 1, 2, and 3 are presumed to originate from three different spectra. Window size *w*_1_ correctly detects four VLM groups. Window size *w*_2_ however is too wide and will detect ambiguous and erroneous groups. Moreover, *w*_2_ will detect several overlapping VLM groups.

Hence, this motivates the following definition of overlapping VLM points. Given (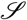, *w*), a VLM *v*_*i*_ w.r.t. (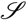, *w*) is said to *overlap* with another VLM *v*_*j*_ (with respect to (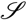, *w*)) if and only if there exists an intersection between the *m*/*z* interval [*v*_*i*_(1 −*w*), *v*_*i*_(1 + *w*)] and the *m*/*z* interval [*v*_*j*_(1 − *w*), *v*_*j*_(1 + *w*)]. Moreover, we say that a VLM *v* w.r.t. (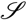, *w*) is *isolated* from all all other VLM with w.r.t. (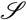, *w*) if and only if there does not exists any other VLM *v*′ w.r.t. (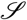, *w*) that overlaps with *v*. For a given window size *w*, the algorithm that we present in the next subsection identifies *all* isolated VLM points w.r.t. (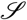, *w*). Consequently, the best value for *w* is one for which the number of isolated VLM points is the largest.

### An Algorithm for Virtual Lock Mass Identification

Given a set 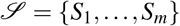 of *m* spectra, each peak is identified by a pair (σ, *ρ*) where σ ∈ {1,…, *m*} is the index of its spectrum of origin and *ρ* ∈ {1,…, *n*_σ_} is the index of the peak in spectrum *S*_σ_ containing *n*_σ_ peaks. Given that we have a total of *n* peaks in 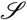, we have that 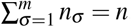. For the description of the algorithm, *μ*(σ, *ρ*) denotes the *m*/*z* value of peak (σ, *ρ*). Finally, we assume that the peaks in each spectra *S*_*i*_ are listed in increasing order of their *m*/*z* values.

The proposed algorithm uses two data structures: a *binary heap* and a so-called *active sequence*. A binary heap is a classical data structure used for priority queues which are useful when one wants to efficiently remove the element of highest priority in a queue. In our case, the heap will maintain, at any time, the next peak of each spectra to be processed by the algorithm. Hence, given a set 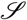 of *m* spectra, the heap generally contains a set of *m* peaks, where each peak belongs to a different spectrum of 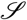. The “priority value” for each peak (σ, *ρ*) in the heap is given by its *m*/*z* value *μ*(σ, *ρ*); a peak with the smaller mass is always on top of the heap. A heap *H* containing the first peak of each spectrum can thus be constructed in *O*(*m*) time. Moreover, we can read the *m*/*z* value at the top of the heap in constant time; we can remove the peak (σ, *ρ*) of the top of the heap and replace it with the next available peak in the spectrum *S*_σ_ in *O*(log*m*) time.

The second data structure is, what we call, the *active sequence A*. At any time, *A* contains a sequence of peaks, listed in increasing order of their *m*/*z* values, which are currently being considered to become a VLM sequence. That data structure uses a doubly linked list *L* and a boolean-valued vector *B* of dimension *m*. The linked list *L* is actually containing the sequence of peaks to be considered for the next VLM and the vector *B* is such that, at any time, *B*[σ] = *True* if and only if a peak from spectrum *S*_σ_ is present in *L*. The active sequence *A* also maintains the *m*/*z* value *μ*_*l*_ of the last peak that was removed from *L*, the average *m*/*z* value *μ*_*A*_ of the peaks in *L*, and a copy *w*_*A*_ of the window size *w* chosen by the user. Since *L* is a linked list, we can read the front (first) and back (last) values of *L* in constant time, as well as obtaining its size (number of peaks). Removing the value at the front of *L* is also performed in constant time.

We now present a short description of the algorithm for virtual lock mass identification. The fully detailed description is provided in Supplementary Information 1.

#### Validation of an active sequence

For this step, we use a method, call *A.isValid*(), that returns *True* if and only if the peaks in the active sequence *A* satisfies all the criteria enumerated in the definition of a VLM. A precondition for the validity of this method is that *L* contains only peaks that belong to distinct spectra of 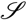. This precondition holds initially for an empty list *L* and will always be maintained for each new peak inserted in *A* (see the next paragraph for details). Thus, this step of the algorithm checks first that the active sequence contains exactly 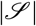 peaks, thus one peak from each spectrum in the set. Then, if there are still peaks in the heap, we verify that the peak at the top of *H* (thus, the peak immediately following the active sequence) has a *m/z* value that is out of the interval [*μ*_*A*_(1 − *w*),*μ*_*A*_(1 + *w*)]. Similarly, it is verified that the peak whose *m*/*z* value immediately precedes the active sequence also has an *m*/*z* value outside of [*μ*_*A*_(1 − *w*),*μ*_*A*_(1 + *w*)]. If either peak lies inside this window, then the property (4) of a VLM is violated, as the window contains more than 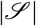 peaks. Finally, we ensure that the first and last peaks in the active sequence *A* are both within the window [*μ*_*A*_(1 − *w*),*μ*_*A*_(1 + *w*)]. If all checks pass, then the current sequence is considered a potential virtual lock mass.

#### Advancing the active sequence

This step tries to insert at the end of the list *L* of *A* the peak (σ, *ρ*) located on top of *H*. The insertion succeeds if the resulting *A* still have some probability that the peak sequence can become a VLM after zero or more future insertions. Thus, we first verify if another peak from spectrum 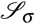 is present in *A*. If that is the case, then the insertion fails. Otherwise, we compute the new value 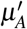 that *μ*_*A*_ will have after the insertion. If the peak at the front of *L* (the peak in *A* having the smallest *m*/*z* value) and the new peak (σ, *ρ*) have masses that are within the window 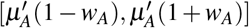, then the insertion succeeds. The peak is inserted, and *H* is updated by removing the peak (σ, *ρ*) and adding the next peak from the spectrum 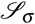. Thus, this step ensures that we can insert a new peak in *A* and still have some probability that the sequence can become a VLM after zero or more future insertions.

Whenever we have an insertion failure, it means that the active sequence cannot become a valid VLM and that we must remove from *A* the peak having the smallest *m*/*z* value (which is located in the front of *L*) in order to have a chance that the sequence of peaks in *A* becomes a valid VLM.

#### Advancing the lower bound

This step is used to remove the peak (σ, *ρ*) at the front of *L* until a valid insertion can be made. First, it updates *B*[σ] to *False*, as peak (σ, *ρ*) is about to be removed and no peak from *S*_σ_ will be in the active sequence *A* anymore. The *m*/*z* value of peak (σ, *ρ*) is copied in *μ*_*l*_, and the peak is then removed from *L*. If *L* is empty at this point, its average *m*/*z* value *μ*_*A*_ is set to 0. Otherwise, *μ*_*A*_ is set to the average value of the peaks remaining in the active sequence.

#### Removing overlapping virtual lock masses

The final step of the algorithm removes all overlapping VLMs. As described in Appendix 1, a Boolean vector (with a number of components equal to the number of VLMs found) is initialized to *False*. Then, we simply iterate over all the VLM points found and assign the corresponding vector entry to *True* whenever a VLM point (with *m*/*z* value *μ*) is found such that its window [*μ*(1 − *w*),*μ*(1 + *w*)] overlaps with that of its neighboring VLMs. Only the VLMs whose entry in the vector is *False* are kept.

#### Main algorithm

Having described the data structures used and their methods, we are now in position to present the main algorithm for virtual lock mass detection, which is described by Algorithm (1). The task of this algorithm is to find all the isolated VLM points w.r.t. (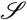, *w*). To achieve this, the central part of the algorithm is to find find the sequence 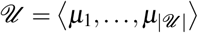 of all possible VLM points w.r.t. (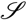, *w*). This sequence may contain several pairs of overlapping VLMs. The strategy to achieve this central task is to use *A.insert*(*H*, 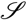) to try to insert in *A* (consequently in *L*) the next unprocessed peak of 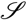, which is always located on the top of the heap *H*.

Initially, the first peak of 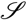, a peak having the smallest *m*/*z* value among those in 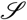, gets eventually inserted in an empty *A* by *A.insert*(*H*, 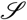). Next, after verifying with *A.isValid*(*H*) if the content of *A* satisfies the criteria to be a valid VLM sequence, we try to insert again in *A* the next available peak. On each insertion failure, we test if, before this insertion, the content of *A* was a valid VLM sequence. This is done with the Boolean variable *f ound* (which is set to *True* as soon as the content of *A* is a valid VLM sequence and which is set to *False* immediately after the average *m*/*z* value *μ*_*A*_ of *A*’s content is appended to 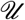). Hence, for each considered peak in *L. f ront*(), we try to insert one more peak in *L* and test after the insertion if *L*’s content is a valid VLM sequence. If we cannot insert an extra peak in *L* with the current peak in *L. f ront*() this means that there is no possibility of finding one more VLM sequence with the current peak in *L. f ront*(). In that case we remove that peak from *L* with *A.advanceLowerBound*() and, consequently, *L. f ront*() now becomes the peak that was next to *L. f ront*() in *L*.

**Algorithm 1:**
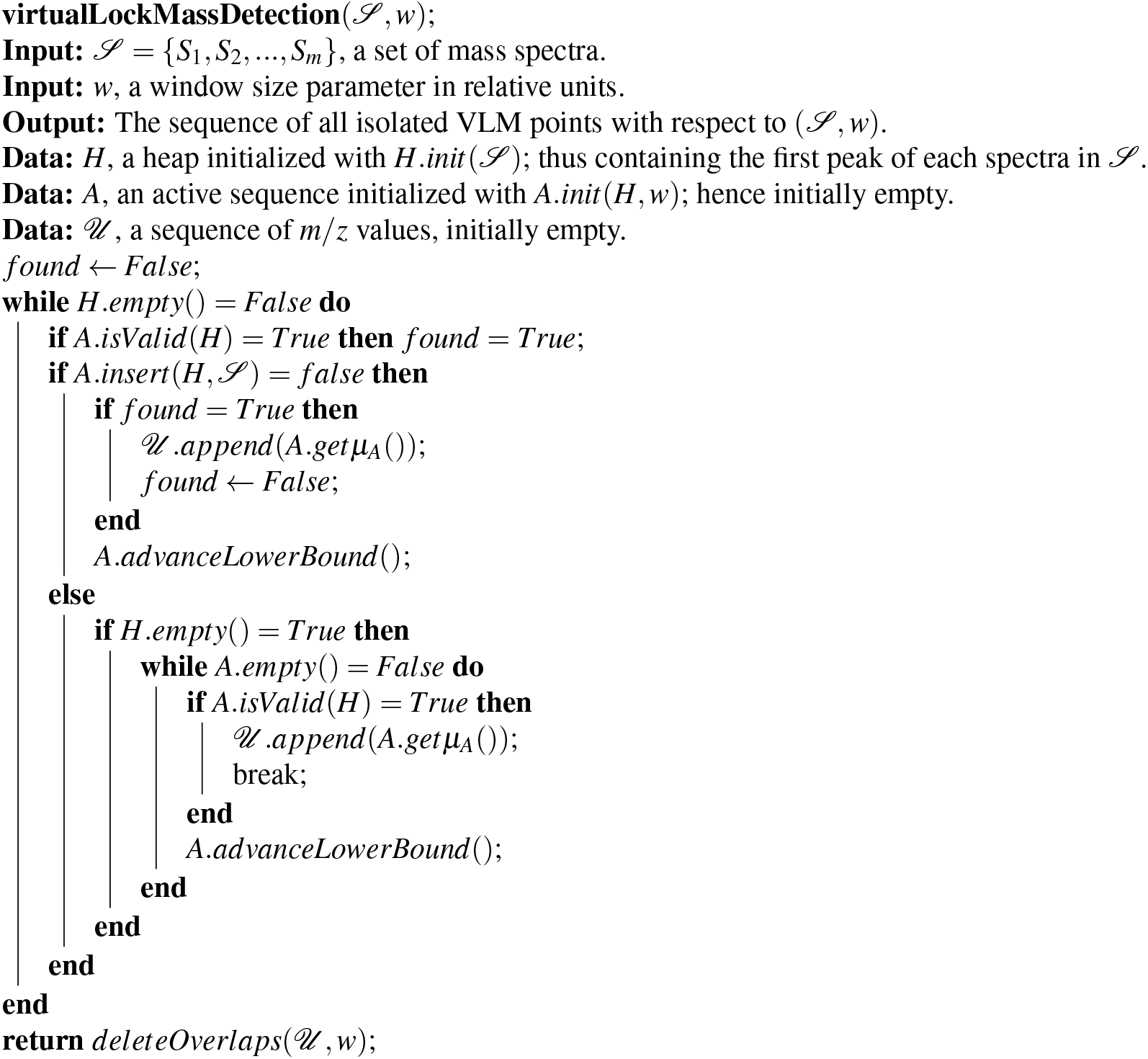
The Virtual Lock Mass Detection Algorithm.

Hence, with this strategy, the algorithm attempts to find the largest consecutive sub-sequence of peaks from 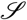 that starts with any given peak in 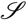 and that forms a valid VLM sequence. In addition, note that in the **else** branch of Algorithm (1), we verify if *H* becomes empty after a successful insertion. In that case, we need to check if we can find a valid VLM sequence by incrementing sequentially the lower bound *L. f ront*() and then append to 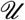 the first VLM found. Then, we can safely exit the **while** loop since any other possible VLM sequence will be a subset of the one already found. Without this **else** branch, a VLM sequence that ends with the last peak presented by *H* would be missed by the algorithm.

As explained in Appendix 1, the running time of Algorithm (1) (i.e., the VLM detection algorithm) is in *O*(*n* log*m*) for a sequence 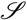 of *m* spectra that contains a total of *n* peaks. This, however, is for a fixed value of window size *w*. Note that in order to obtain the most accurate correction (by interpolation) for the spectra in 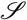, we should use the largest number of isolated (i.e., non-overlapping) VLMs we can find. Consequently, the optimal value for *w* is the one for which Algorithm (1) will give the largest number of isolated VLMs. Moreover, note that if *w* is too small, very few VLMs will be detected as *w* will not be able to cover exactly one peak per spectra. If, on the other hand, *w* is too large, a large number of the VLMs found in the first phase of the algorithm will overlap and the remaining isolated VLMs will be rare. Consequently, because of this “unimodal” behavior, one can generally find rapidly the best value for *w*. In our case, we never needed to tried more than 20 different values.

### An Algorithm for Virtual Lock Mass Correction

Given a set 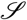 of spectra and a widow size parameter *w* expressed in relative units, once the sequence 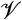 of all isolated VLM points w.r.t. (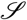, *w*) has been determined, the individual spectra in 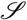 can be corrected in a manner similar as it is usually done with traditional lock masses. Algorithm (2) performs the correction needed for each peak in a spectrum 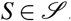.

**Algorithm 2:**
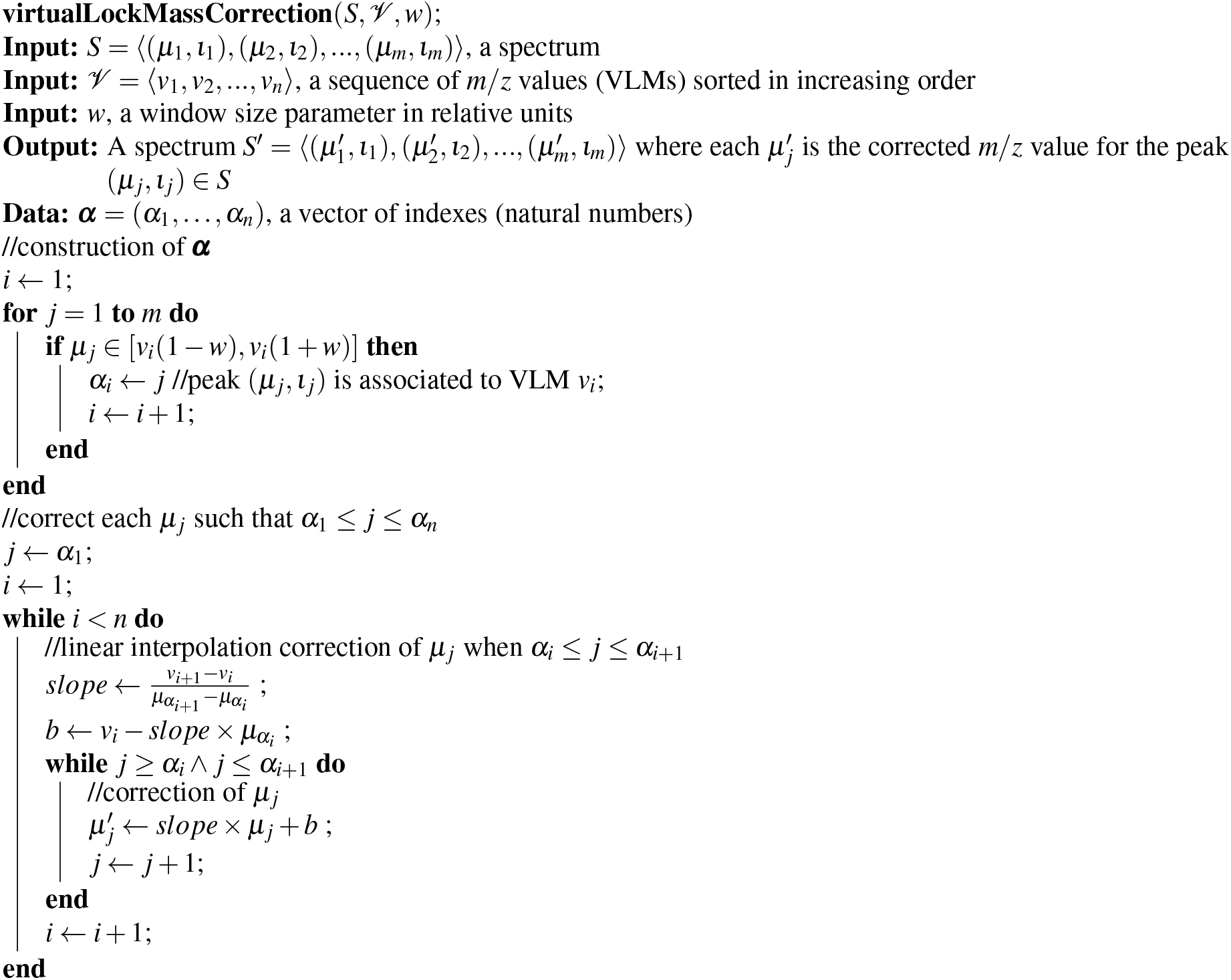
Virtual Lock Mass Correction Algorithm

First, in the **for** loop, we identify each peak of *S* corresponding to a lock mass point 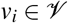. Since 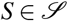 and *v*_*i*_ is a VLM point w.r.t. (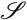, *w*), we are assured to find exactly one such peak *p*_*j*_ ∈ *S* with an observed *m*/*z* value of *μ*_*j*_ such that *μ*_*j*_ lies in the interval [(1 − *w*)*v*_*i*_, (1 + *w*)*v*_*i*_]. For such *μ*_*j*_, we assign the index *j* to *α*_*i*_ so that ***α*** = (*α*_1_,…, *α*_*n*_) is a vector of *n* indexes, each pointing to the peak in *S* associated to a VLM point. Note that for *μ*_*j*_ ∈ [(1 − *w*)*v*_*i*_, (1 + *w*)*v*_*i*_], its corrected *m*/*z* value must be equal to *v*_*i*_. Instead of performing these corrections immediately in the **for** loop, we delay them to the linear interpolation step where all peaks having a *m*/*z* value *μ*_*j*_ such that *α*_1_ ≤ *j* ≤ *α*_*n*_ will be corrected.

Next, for each VLM *v*_*i*_, we correct by linear interpolation all the *m*/*z* values *μ*_*j*_ such that *α*_*i*_ ≤ *j* ≤ *α*_*i*+1_. To explain precisely this procedure, let *μ*′(*μ*_*j*_) denote the corrected value of *μ*_*j*_. Linear interpolation consists at looking for a correction of the form

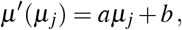

where *a* is called the *slope* and *b* is the *intercept*. By imposing that *μ*′(*μ*_*j*_) = *v*_*i*_ for *j* = *α*_*i*_ and *μ*′(*μ*_*j*_) = *v*_*i*+1_ for *j* = *α*_*i*+1_, we find that

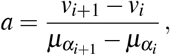

and 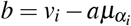. The nested **while** loops of the algorithm performs exactly these linear interpolation corrections for all *μ*_*j*_ such that *α*_*i*_ ≤ *j* ≤ *α*_*i*+1_ for *i* = 1 to *n* − 1.

Once all *m*/*z* values *μ*_*j*_ such that *α*_1_ ≤ *j* ≤ *α*_*n*_ have been corrected, the algorithm is done. Hence, we have decided not to correct any *m*/*z* value of *S* that is either smaller that *v*_1_(1 − *w*) or larger than *v*_*n*_(1 + *w*) because such a peak has only one adjacent VLM and, consequently, could only be corrected by extrapolation, which is much less reliable than interpolation. Therefore, we recommend removing all these peaks from *S* to perform statistical analyses or machine learning experiments. Finally, the intensities of the peaks remain unchanged. The running time complexity of this algorithm is *O*(*m*) where *m* is the number of peaks in the spectrum *S* (see the full details in Appendix 1).

### From VLM correction to spectra alignment

After running the VLM detection and correction algorithms, all the peaks associated with VLM points will be perfectly aligned in the sense that each peak in different spectra associated to a VLM point *v* will have exactly the same *m*/*z* value *v*. However, all the other peaks corrected by Algorithm (2) will not be perfectly aligned in the sense that a molecule fragment responsible for a peak in different spectra will not yield exactly the same mass after correction. This is due to possibly many uncontrollable phenomena that vary each time a sample gets processed by a mass spectrometer, and by the fact that the correction of each peak was performed by an approximate numerical interpolation. However, if all the peaks have been corrected by Algorithm (2), we expect that the peaks corresponding to the same molecule fragment *f* across different spectra will have very similar masses and will all be localized within a very small window of *m*/*z* values. Moreover, we also expect that the *m*/*z* values of the peaks coming from another molecule fragment *g* having a different mass will not cross the *m*/*z* values coming from molecule fragment *f*.

More precisely, suppose that we have executed Algorithms (1) and (2) with a window size parameter *w* (in relative units) on a set 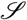 of mass spectra. In addition, suppose that a molecule fragment *f* gives rise to a peak of *m*/*z* value *μ*_1_ in spectrum *S*_1_, and a peak of *m*/*z* value *μ*_2_ in spectrum *S*_2_, and so on for a sub-sequence of spectra in 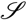. Let 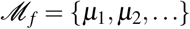 be the set of these *m*/*z* values. Moreover, let *μ*_*f*_ be the average of the *m*/*z* values in 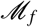. Then, we expect that there exists a window size *θ* in relative units, such that 0 < *θ* ≪ *w*, and for which we have *μ*_*i*_ ∈ [*μ*_*f*_(1 − *θ*), *μ*_*f*_(1 + *θ*)] for all 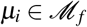. Moreover, if *θ* is sufficiently small, we expect that the sequence 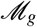 referring to peaks produced by another molecule fragment *g* having a different mass will be such that each 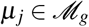 will not be located within [*μ*_*f*_(1 − *θ*), *μ*_*f*_(1 + *θ*)].

Motivated by this hypothesis, let us introduce the following definitions. Given that Algorithms (1) and (2) have been executed on a set 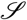 of mass spectra with window size parameter *w* in relative units, and given that we have another window size parameter *θ* ≪ *w* in relative units, we say that a *m*/*z* value *μ*_*f*_ is an *alignment point* w.r.t. (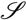, *θ*) if there exists a set 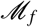 of peaks from 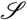 that satisfies the following properties.

1. Every peak in 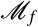 comes from a different spectrum of 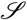.
2. The average of the *m*/*z* values of the peaks in 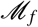 is equal to *μ*_*f*_.
3. Every peak in 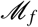 has an *m*/*z* value in [*μ*_*f*_(1 − *θ*), *μ*_*f*_(1 + *θ*)] and all other peaks of 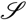 have an *m*/*z* value outside this interval.
4. There does not exist another peak in 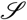 that we can add to 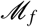 and still satisfy the above properties.

Whenever these criteria are satisfied, we say that 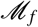 is the *alignment set associated* to alignment point *μ*_*f*_. Given 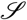 and *θ*, an alignment point *μ*_*f*_ w.r.t. (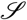, *θ*) is said to *overlap* with another alignment point *μ*_*g*_ w.r.t. (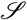, *θ*) if and only if there exists a non-empty intersection between the *m*/*z* intervals [*μ*_*f*_(1 − *θ*), *μ*_*f*_(1 + *θ*)] and [*μ*_*g*_(1 − *θ*), *μ*_*g*_(1 + *θ*)].

Let 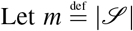. Note that there are only two differences between the definition of alignment point (and its associated alignment set) and the definition of VLM point (and its associated VLM set). The first difference is the fact that a VLM set must contain exactly *m* peaks, whereas an alignment set can contain any number of peaks between 1 to *m* (since the peaks in an alignment set may originate from a molecule fragment which is not present in all the samples for which we have a spectrum in 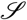). Hence, if we remove the constraint that each virtual lock mass must be formed of 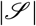 peaks from the validation step, Algorithm (1) then finds all the maximum-length sub-sequence of peaks that satisfy the 4 criteria for a valid alignment set when it reaches the overlap deletion step. The second difference is that there is no intensity thresholds *t*_*a*_ and *t*_*b*_ applied to the peaks for alignment, as we wish to align every peak in the spectra if possible. Note that, generally, a lower intensity threshold is still applied to the peaks in order to remove peaks that are the result of background noise. Consequently, with that very minor change,

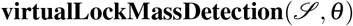

finds all isolated alignment points w.r.t. (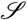, *θ*) in *O*(*n* log*m*) time, where *n* is the total number of peaks in 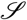.

If the window size parameter *θ* is too large, then many alignment points will overlap and Algorithm (1) will return very few isolated alignment points. If *θ* is too small, then, in contrast with the VLM identification case, Algorithm (1) will return a very large number of isolated alignment points associated to aligned sets that contain only one point. Hence, in contrast with the VLM identification case, the best parameter *θ* is not the one for which we obtain the largest number of alignment points.

What should then be the choice for *θ* ? To answer this question, we consider each VLM point (and its associated sequence of peaks) found by Algorithm (1). If we leave out one VLM point *v*_*i*_ from the correction algorithm (2) and use this algorithm to correct all the *m*/*z* values of the peaks associated to this VLM point, the maximum deviation from *v*_*i*_ among these *m*/*z* values will give us the smallest window size *θ*_*i*_ such that each *m*/*z* value will be located within [*v*_*i*_(1 − *θ*_*i*_), *v*_*i*_(1 + *θ*_*i*_)]. Essentially, this window size *θ*_*i*_ is the smallest one for which we can still recognize all the peaks associated to the same VLM *v*_*i*_. It would then certainly be a very good choice for *θ* in that region of *m*/*z* values. We can then repeat this procedure for all isolated VLM points (except the VLMs with the smallest and largest *m*/*z* values) found by Algorithm (1) to obtain a sequence of *θ*_*i*_ values.

One interesting possibility for *θ* is the maximum among the *θ*_*i*_ values. However, this is clearly an overestimate of the maximum spreading of peaks associated to the same molecule fragment since all the VLMs will be used for the correction, including the one that was left out. Moreover, as we can see in Figure (2), we can recover a large fraction of the non-overlapping VLMs if we use a significantly smaller window size than the max_*i*_*θ*_*i*_. For that reason, we have decided to use, for the window size *θ*, the smallest value covering 95% of the non-overlapping VLMs, i.e., the 95th percentile. Alternatively, to attempt to maximize the accuracy of a learning algorithm, a percentile *z* can be selected by cross-validation along with the selection of the hyperparameters of the learning algorithm.

**Figure 2.**
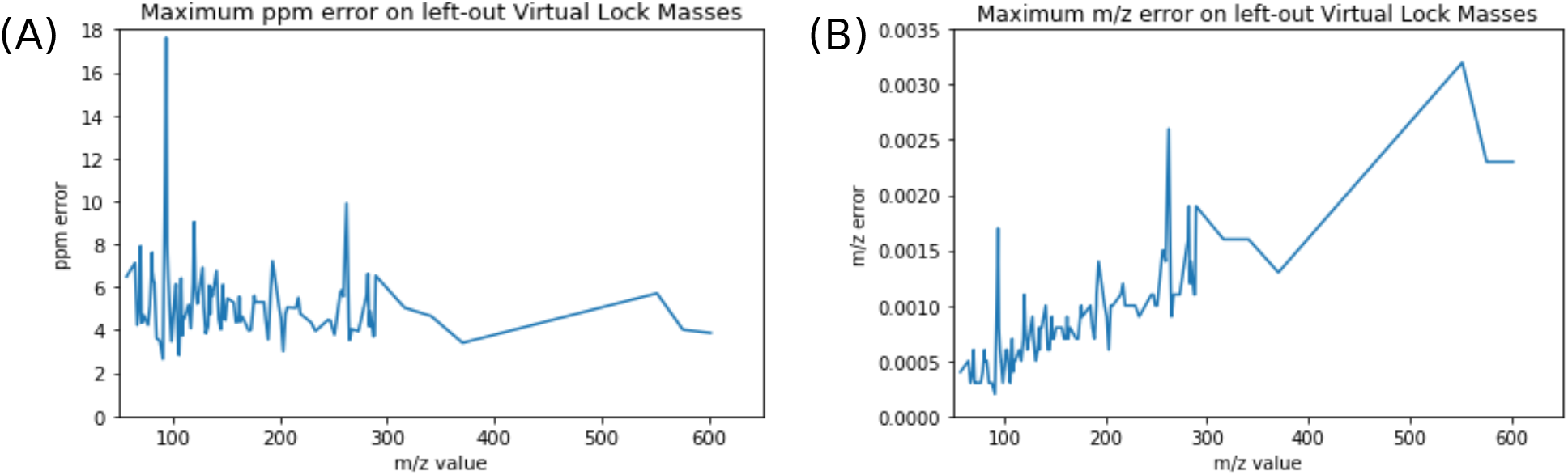
Error in ppm versus mass units. Subfigure (A) shows the error on left-out VLMs in ppms, while Subfigure (B) shows the error in Daltons. This data was acquired on the Days Dataset.

If we have *r* VLM points, each *θ*_*i*_ associated to the *i*th VLM point is found in *O*(*m*) time for a sequence 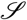 of *m* spectra; thus implying a running time in *O*(*mr*) to find every *θ*_*i*_. Then, the 95th percentile is found by sorting the vector of *θ*_*i*_s in *O*(*r* log*r*) time. Assuming that we always have log(*r*) < *m*, the total running time to find *θ* is in *O*(*mr*), and hence in *O*(*n*) when 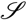 contains a total of *n* peaks. Once the window size *θ* is found, we can then run Algorithm (1) just once on the full set 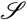 of spectra with that value of *θ* in *O*(*n* log*m*) time. Consequently, the total running time of the alignment algorithm, which includes the running time to find *θ* and to find all non overlapping alignment points w.r.t. (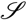, *θ*), is in *O*(*n* log*m*).

Once we have the VLM points and the alignment points, these are used to provide a *representation* of the spectra which is well suited for running machine learning algorithms on them. Indeed, consider Fig (3). For any new spectrum *S*, the VLM points are first used correct the *m*/*z* value of each peak of *S* and, following that, the intensity of any corrected peak that fall into the window associated to an alignment point give a feature of *S*. Hence, the vector of these intensities provides a new representation of the spectrum *S* that we will use for the input into a classifier to predict the label of *S*.

**Figure 3.**
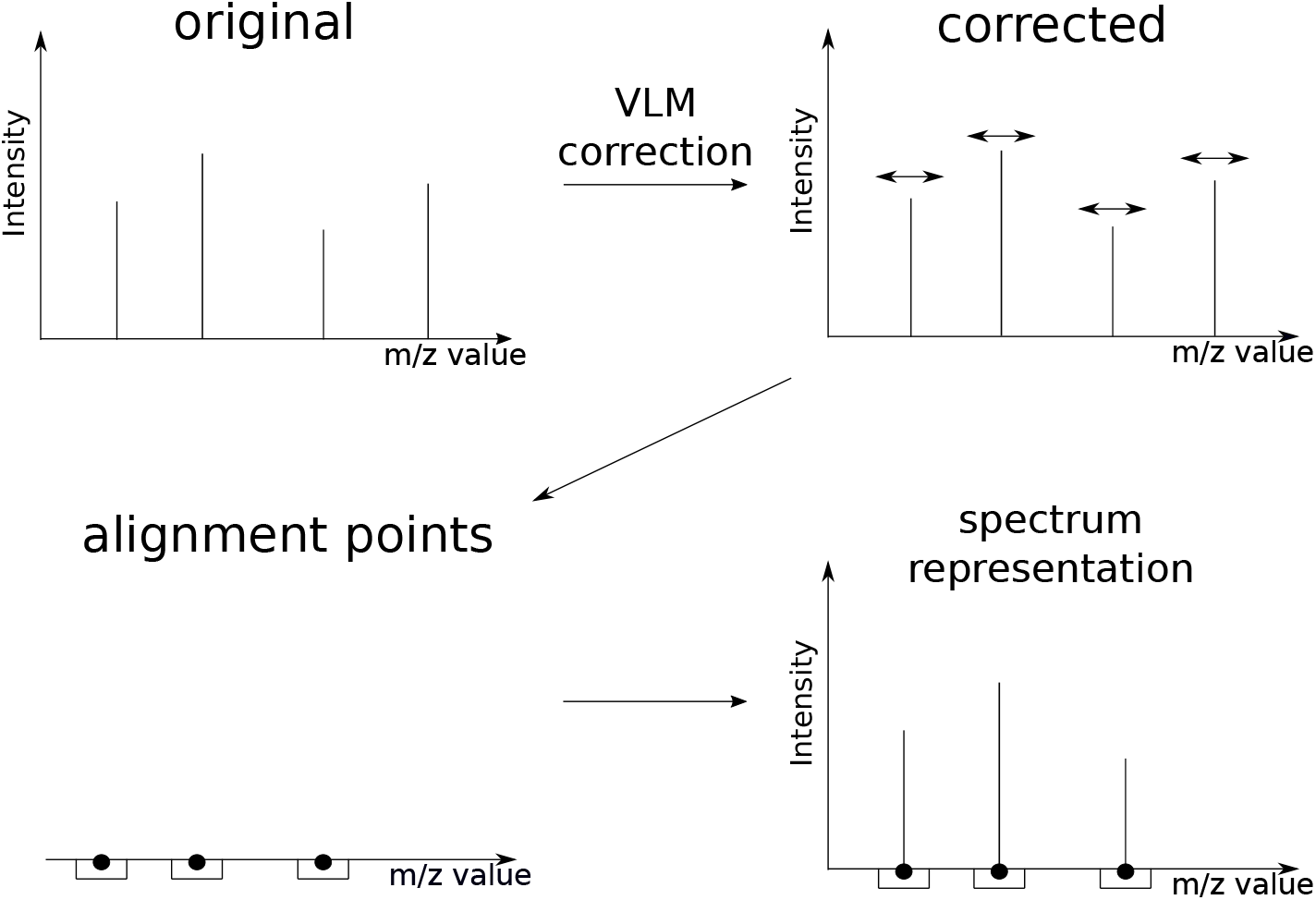
Workflow of the VLM and alignment algorithms. First, VLM points are detected in the original spectra in the dataset and VLM correction is applied. The alignment algorithm is then applied to the corrected spectra in order to obtain the alignment points. The representation of a given spectrum is the subset of peaks which fall within a mass window of an alignment point, with unmodified intensity.

Finally, it might be tempting to use a clustering approach to solve the problem of finding the isolated alignment points. However, we have to keep in mind that current trends lead to the processing of hundreds of spectra, each potentially containing thousands of peaks. The total number of peaks to be processed can thus reach a million peaks or more. In our case, the total running time of the full pipeline (finding all isolated VLMs, correcting all the mass spectra with the VLMs, and finding all the isolated alignment points) is in *O*(*n* log*m*). Hence, any algorithm running in Ω(*n*^2^) time, will be completely surpassed by the proposed pipeline of algorithms. Currently, the running times of popular off-the-shelf clustering algorithms such as K-means and linkage-based clustering algorithms all require a running time in Ω(*n*^2^). Moreover, all the clustering algorithms that we know have at least one parameter to tune, which often includes the number of desired clusters. Hence, with the current state of knowledge, a clustering-based algorithm is bound to be substantially less efficient than the proposed pipeline of algorithms.

An implementation of the algorithms for Python is available at https://github.com/francisbrochu/msvlm.

### Dataset descriptions

#### *Days* Dataset

Plasma from 20 healthy individuals was equally pooled together. The pooled plasma was aliquoted and kept at −20 °C.

On two consecutive days, an acetonitrile crash was performed using 9 parts of acetonitrile (Fisher Optima) for 1 part of unfrozen plasma pool. The crash solution was centrifuged at 4000 rpm for 5 minutes. 2 *μ*l of the solution was spotted on every well of a 96 wells Lazwell plate (Phytronix). The same experiment was repeated the next day. The dataset is then formed of the 96 aliquots unfrozen on the first day and the 96 aliquots of the second day, forming a 192 samples dataset.

#### *Clomiphene-Acetaminophen* Dataset

One pill of acetaminophen (500 mg) was diluted in 50 ml of methanol and water (50:50). The solution was put in a sonicating bath for 20 minutes. The resulting solution was centrifuged and diluted 1:100 in water. For clomiphene, we used a solution of 100*μ*g/ml of clomiphene in methanol (Phytronix).

A pool of plasma was crashed as previously described. The solution was split in 3 parts. One received 10 *μ*l of the acetaminophen solution, another 10 *μ*l of the clomiphene solution and the last one stayed unmodified. Each type of sample was spotted 32 times.

#### *Malaria* Dataset

*Plasmodium falciparum* parasites were put in culture in red blood cells and tightly synchronized. Culture was performed for 28-36 hours, until parasites are in the trophozoite stage and parasitaemia reached 5-10%. In the same conditions, red blood cells were kept uninfected. Cells were diluted to 2% hematrocrit by adding the correct amount of pelleted cells to complete RPMI media. 200 *μ*l of the cell suspensions was deposited in a 96 well plate in order to have 40 samples of infected cells and 40 samples of uninfected cells.

After 4 hours at 37 °C, the plate was spin at 800x *g* for 5 minutes. 180 *μ*l of culture media was removed. Pellet was resuspended in the remaining 20 *μ*l and 10 *μ*l was transferred to a new 96 well plate. 100 *μ*l of ice-cold methanol was quickly added to the plate and put on dry-ice to stop any metabolic reaction. The plate was vortexed 3 times, for 15 seconds each, over 15 minutes incubation on dry-ice. The plate then was placed for sonication in a water bath for 5 times 1 minute with 2 minutes breaks on dry-ice. Finally, the plate was centrifuged at 3200 rpm for 5 minutes at 4°C. 30*μ*L of the supernatant was transferred to another plate and kept at −80°C until LDTD-MS analysis. For analysis, 2 *μ*l of the metabolomic extract was spotted on a 96 well Lazwell plate and left at room temperature until dryness.

#### *Cancer* Dataset

Plasma from patients diagnosed for breast cancer and from healthy patients were individually treated using the same acetonitrile crash protocol. A total of 96 samples from breast cancer patients were acquired. In addition, 96 plasma samples from healthy patients were also acquired in order to have control samples.

##### Data acquisition

All data were acquired on a Synapt G2-Si mass spectrometer. The instrument was operated in high resolution mode. Except if stated otherwise, data acquisition was performed in positive ionization. The acquisition method was *MS*^*e*^ with a scan time of 0.1 second. Calibration of the instrument was performed daily before data acquisition using a solution of sodium formate 0.5 mM. The instrument was operated with Mass Lynx software. The source is a LDTD 960 ion source (Phytronix). The laser pattern used is the following: 2 seconds at 0%, ramp up to 65% in 6 seconds, hold at 65% for 2 seconds and back at 0% in 0.1 second.

#### Data conversion

Raw files produced by the mass spectrometer were converted to ion list using a continuous to centroid approach using the ProcessKernel software (Waters Corporation) using only the first function (low energy) present in the files. The resulting centroided peak list were used for data analysis.

For all experiments presented in this article, the *t*_*a*_ threshold on intensity for virtual lock mass detection was set at 1000 counts. No *t*_*b*_ threshold was applied in the experiments since there was no saturation effect detected on the spectra in the datasets.

#### Use of human participants

All participants provided written informed consent, and the study protocol was reviewed and approved by the Research ethics committee of the CHU de Québec-Université Laval Research Center. All experiments were performed in accordance with relevant guidelines and regulations.

## Results

### A consistent set of virtual lock masses can be detected in different batches

This experiment was conducted on the *Days* dataset (see *Methods*), which consist of 192 samples of pooled blood plasma. Half of the samples were acquired on a given day and the others were acquired the next day. Since the samples are of the same nature (as in, they are all of the same type of biofluid), we expect a high similarity apart from inter-batch variations. The goal of the experiment was to determine if a consistent set of virtual lock masses could be detected among similar datasets and within parts of the same dataset.

The VLM detection algorithm was independently applied to 1) every spectrum in the dataset, 2) only the spectra acquired on the first day, and 3) only the spectra acquired on the second day. The algorithm was applied with the same window size of 40 ppm in all cases. This window size was determined by the procedure described in the Methods section, being the *w* that yielded the largest number of isolated VLMs on the entire dataset.

The detected VLMs were then compared in the following manner. We define that if we have two sets of spectra *A* and *B*, their detected VLMs will be 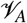 and 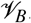. Each element of 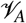 is a VLM *v*_*A*_ that is composed of a single peak per spectrum for the spectra in *A*. If *B* ⊂ *A*, then a VLM 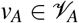 and a VLM 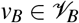 are homologous if the peaks forming *v*_*B*_ are a subset of the peaks forming *v*_*A*_. Additionally, we can define comparisons between the VLMs of subsets of *A*. If we have sets of spectra *A*,*B* and *C*, where *B* ⊂ *A* and *C* ⊂ *A*, then we can define that VLMs *v*_*B*_ and *v*_*C*_ are homologous if *v*_*B*_ if homologous to *v*_*A*_ and *v*_*C*_ is homologous to *v*_*A*_.

We compared the peak groups forming the VLMs in all spectra with the spectra acquired on the first day, and found that the 113 VLMs detected on all spectra have homologues in the set of 148 VLMs detected on the first day. Conversely, we observed that the 113 VLMs also have homologues within the set of 118 VLMs detected in the spectra acquired on the second day.

Hence, the algorithm finds common VLM points in all settings, corresponding to different days and multiple instrument recalibrations. This suggests that it correctly identifies landmark compounds that are present in a particular type of sample, which can be used as a common basis for correction. We therefore conclude that our detection algorithm behaves as expected.

### Virtual lock mass correction improves machine learning analysis

Machine learning experiments were conducted on four binary classification tasks. The first two tasks consist of the detection of a single compound spiked in blood plasma samples from the *Clomiphene-Acetaminophen* dataset (see *Methods*). The third task is the detection of malaria infection in red blood cell culture samples from the *Malaria* dataset (see *Methods*). The fourth and final task consists of distinguishing plasma samples of patients with and without breast cancer in the *Cancer* dataset (see *Methods*).

### Influence of the number of samples on virtual lock mass correction

An experiment was performed in order to evaluate the behavior of the VLM detection and correction algorithms on varying numbers of samples. In a first step, the VLM detection algorithm followed by the VLM correction algorithm was performed on the whole set of spectra. 25 spectra were randomly selected as a test set. These test spectra will be considered the “ground truth”, i.e., the best correction that the algorithm can achieve for these 25 spectra.

The algorithm was subsequently applied to a part of the training set. This part was gradually increased from 10 to 160 spectra. At each point, the uncorrected test spectra were corrected and compared to the ground truth. The difference in *m*/*z* value between the homologous peaks is calculated in ppm. Then, the difference is squared and summed for all test spectra. Finally, this sum is divided by the number of peaks in the test spectra and the square root is taken. The difference in correction is thus expressed as the Root Mean Squared Error (RMSE) in ppm units for each peak. This experiment was repeated 50 times, with randomly re-partitioned test sets, in order to obtain statistically significant results.

Fig 4 shows the learning curves obtained on three different datasets. In each case, the trends is similar. When sub-sampling a low number of spectra as a training set for the VLM detection and correction algorithms, a higher number of lock masses is found. As the number of training spectra increases, the number of virtual lock masses found diminishes and starts to plateau near the number of lock masses found in the whole dataset. This is explained by the fact that when few spectra are in the training set, there is a higher number of candidates. As new spectra are added in the training set, there is a probability that one of the new spectra are missing at least one peak that was previously considered a virtual lock mass. These peaks could be missing because of strong noise, either on the *m*/*z* axis or in terms of intensity, rendering its intensity too small to be considered a VLM. A peak could also be missing simply because the compound or fragment generating that peak is not present in all samples.

**Figure 4.**
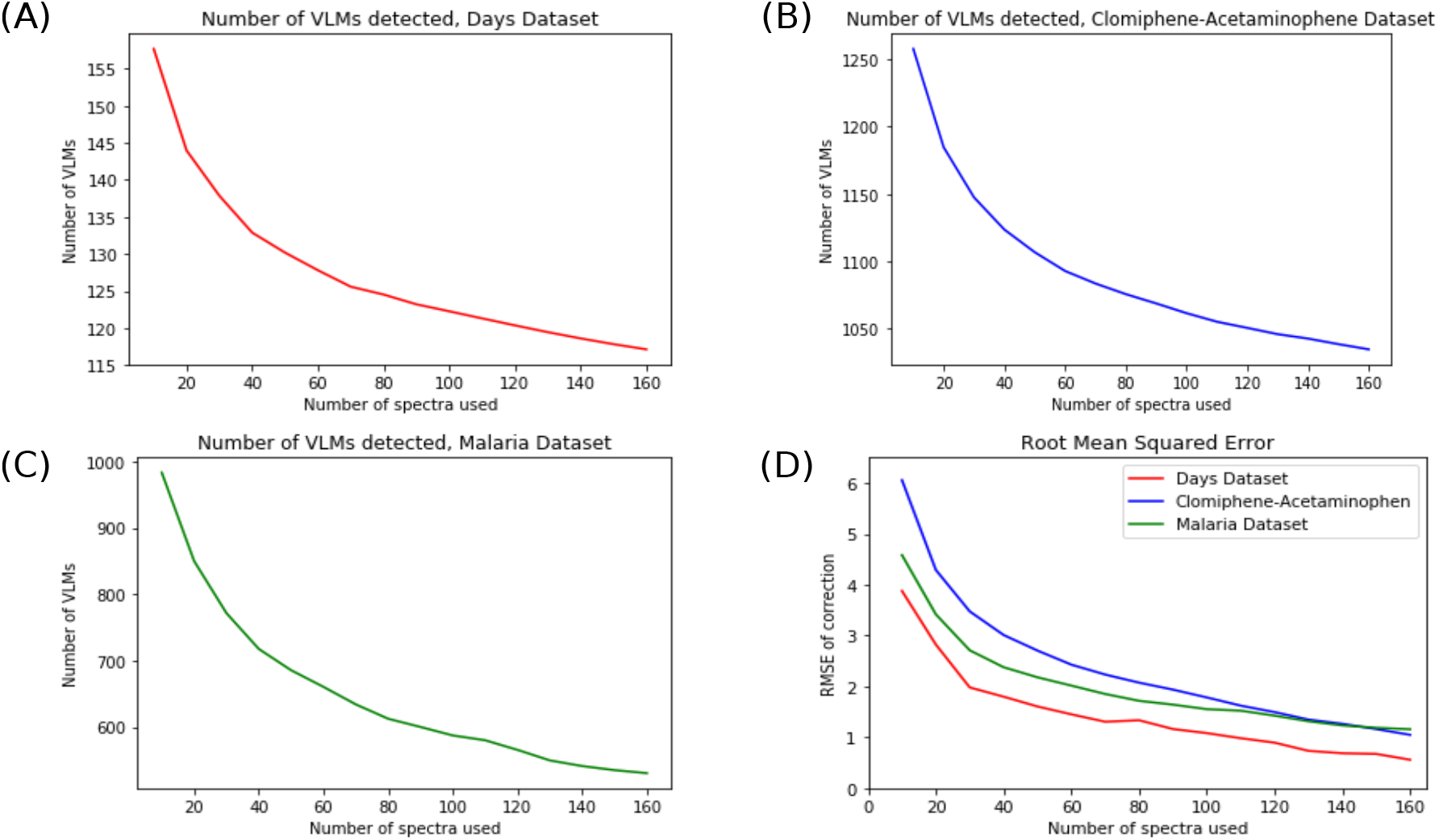
Learning Curves of Virtual Lock Mass Detection and Correction. Subfigures (A), (B) and (C) show the learning curves for three different datasets ((A) Days, (B) Clomiphene-Acetaminophen and (C) Malaria). Subfigure (D) shows the Root Mean Square Error (RMSE) of VLM Correction for these datasets on an unseen test set. This test set consisted of 25 randomly selected samples from the datasets, which were kept separate. The experiments were replicated 50 times and averaged.

The same trend is found in all three datasets for the Root Mean Squared Error (RMSE) in Subfigure (D). The error is initially high when few spectra are in the training set, but as more spectra are added in the training set it gradually decreases. In the case of the *Days* Dataset, the final average RMSE when using 160 spectra to train the algorithm is 0.56 ppms. For the other two datasets (*Clomiphene-Acetaminophen* and *Malaria*), the final RMSEs are approximately 1.10 ppms. In each case, the RMSE drops under 2.0 ppms when using 100 spectra or more to train the correction algorithm. In conjunction with the results of inductive learning shown above, these results suggest that the VLM detection and correction algorithms can generalize the virtual lock masses and correction it learns to unseen spectra of the same nature, such as those of a new test set.

Figure (5) shows boxplots of the RMSE of the peaks at different points in the learning curve. Each subfigure shows the RMSE for peaks found in different mass ranges. In every range, the RMSE diminishes with the added spectra to the training set. Moreover, in addition to the average and median values decreasing, we can observe that the interquantile range also decreases. The outliers also tend to be of lesser values. Where multiple outliers have RMSE’s greater than 15 when correcting based on 10 spectra, the highest values tend to be less than 10 to 12 (depending on the mass range) when correcting based on 150 spectra.

**Figure 5.**
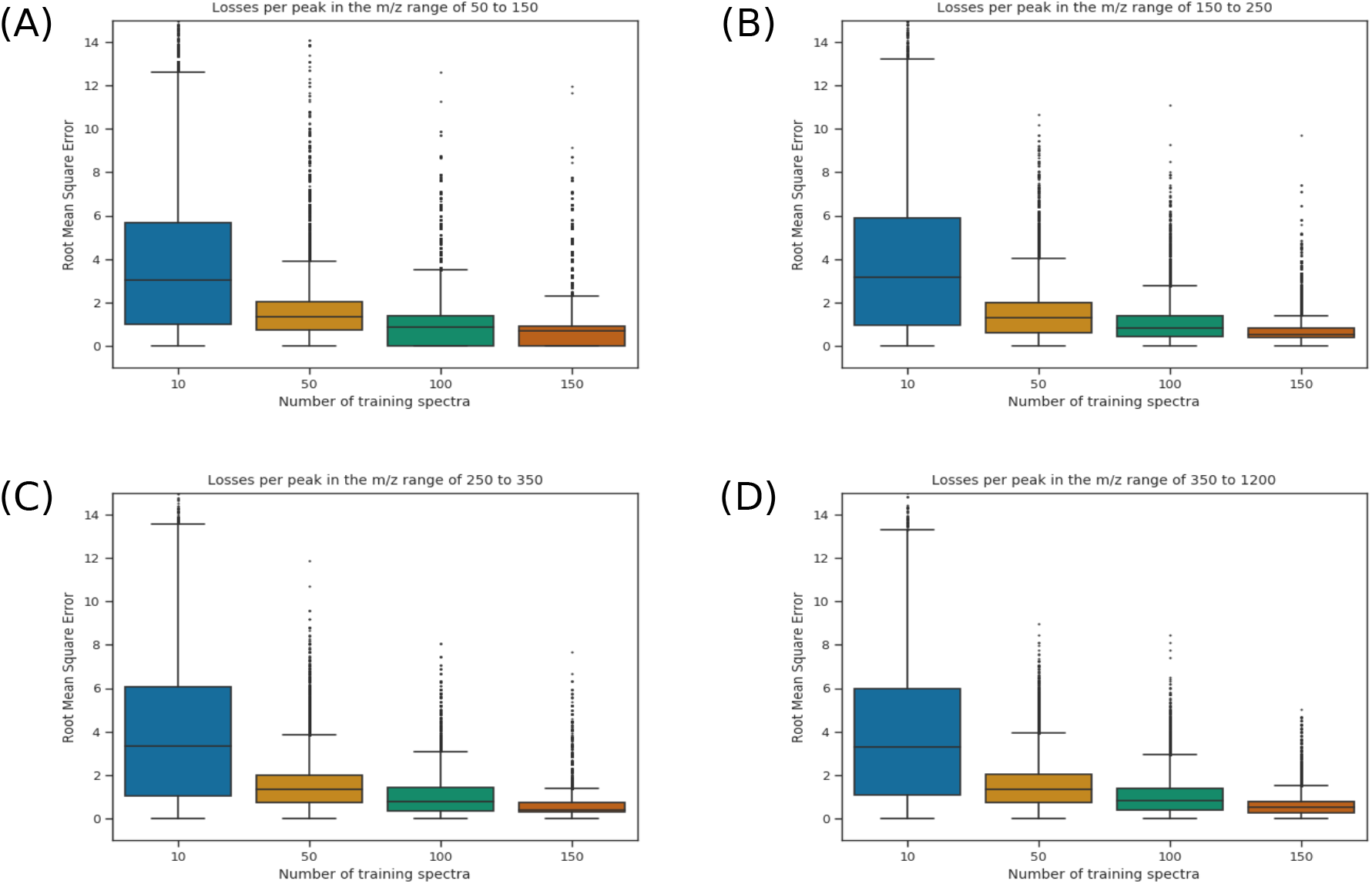
Loss per peak in different *m*/*z* ranges of the spectra. Each boxplot represents the RMSE of the peaks in a given region (50-150 in (A), 150-250 in (B), 250-350 in (C) and greater than 350 in (D)). Shown here are the results for the Days Dataset, in increasing order to training spectra, from 10 to 150. The outliers are shown as ticks over each box.

### Machine learning algorithms

Multiple machine learning algorithms were applied to the spectra. The first algorithm used is the AdaBoost ensemble method^20^. This method learns a weighted majority vote on a set of simple pre-defined classifiers. A linear Support Vector Machine (SVM)^21^ was used with a L1-norm regularizer. The latter is to ensure that the predictions are based on a small subset of the peaks. We also used decision tree^22^ and Set Covering Machine classifiers^23^. These algorithms have the advantage of producing interpretable classifiers that consist of a very small combination of simple rules on peak intensities. We used the Scikit-learn implementations for Python of the AdaBoost, CART, and the L1-regularized SVM^24^ algorithms; whereas we used our own Python implementation of the SCM. This implementation is available at https://github.com/aldro61/pyscm.

### Experimental protocol

For each experiment, the spectra were randomly partitioned into a training set and a test set. For the compound detection tasks (clomiphene and acetaminophen), the test set consisted of 50 selected samples. For the cancer detection task, the same number of samples were included in the test set. Finally, for the malaria detection task, 100 samples were selected for the test set. The hyper-parameters of each learning algorithm were chosen by 5-fold cross-validation on the training set (refer to Elements of Statistical Learning^25^). Each experiment was repeated 10 times independently on different partitions of the data.

Two different experimental protocols were tested which are illustrated in Fig (6). First, the correction and alignment algorithms were applied in the transductive learning setting^26^. In this setting, the whole dataset is exposed to the pipeline of proposed algorithms (VLM detection + VLM correction + alignment point detection). The training and testing sets are then partitioned randomly. The second experimental protocol was conducted as the inductive learning setting, in which the pipeline of proposed algorithms were only applied to the training set. Hence the set of alignment points is found from the training set only. For the inductive learning protocol, the percentile parameter of the alignment algorithm is considered an hyper-parameter and is thus cross-validated on the training set. For the transductive learning protocol, the percentile parameter is set at 95%. The features shown to the machine learning algorithms are the alignment points and their associated intensity values.

**Figure 6.**
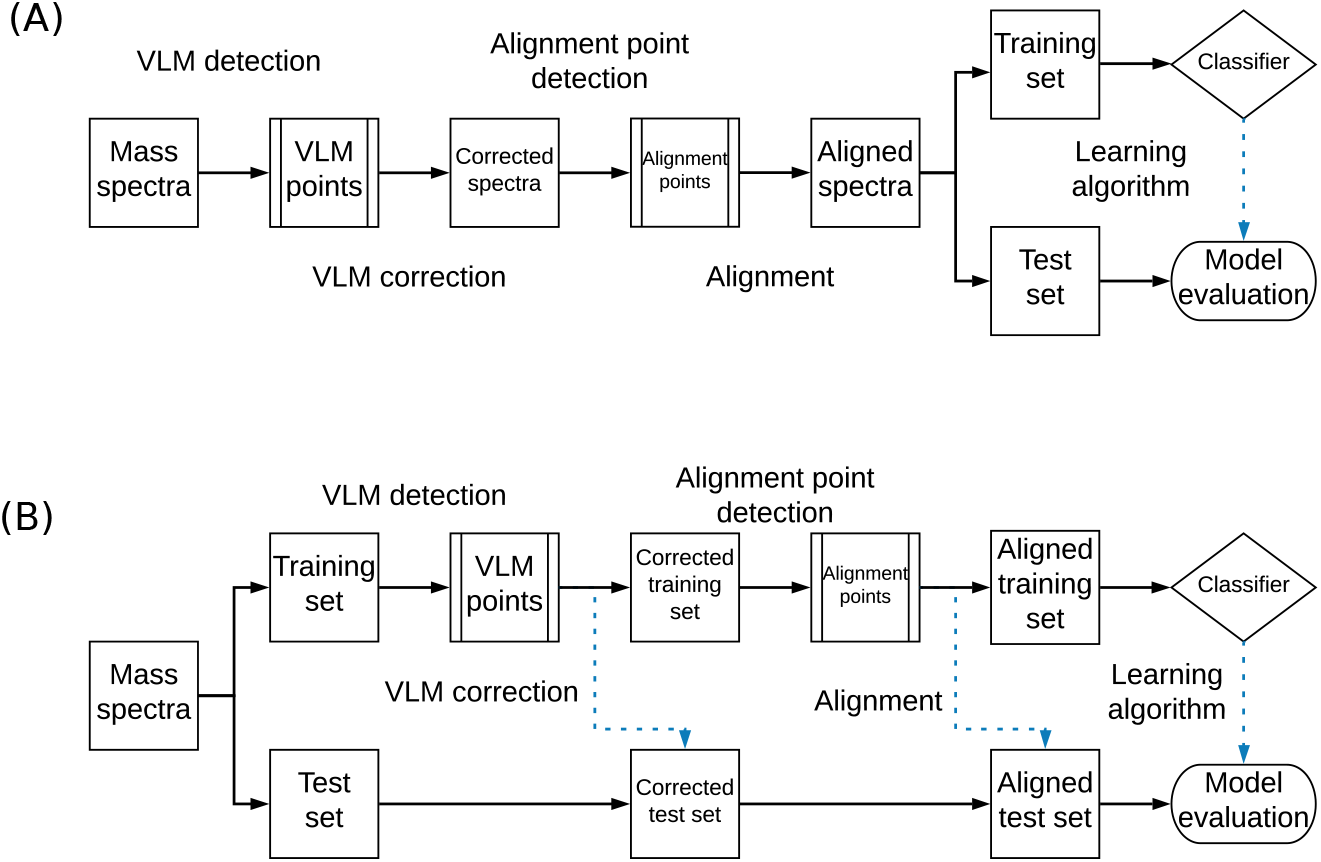
Transductive and inductive workflows. A) The transductive workflow, in which all spectra are corrected at once, prior to partitioning the data into a training and testing set. (B) The inductive workflow, where the data are first partitioned and only the spectra in the training set are used to learn a transformation that is applied to all spectra. The dotted blue arrows show where the algorithms were applied on unseen data, while the whole black arrows show the workflow of the training data. Thus, in the inductive workflow, the test set is formed of unseen data that is only used for the final evaluation of the model. In the transductive case, some information is taken from all samples, while only the learning part of the workflow separating a test set on which the algorithm does not learn.

For each task, we compared the performance of classifiers according to their preprocessing. We thus compared (a) simply binning the spectra, (b) using the VLM detection and correction algorithms and then binning the mass spectra and (c) using the VLM detection and correction algorithms before using the alignment algorithm. Binning is a commonly used technique in mass spectrometry analysis consisting in grouping peaks and intensities found in a larger bin on the *m*/*z* axis into a single point or peak^27^.

### Results for transductive learning

Table 1 shows the results of the machine learning experiments in the transductive setting for different tasks. Let us first consider the case of the clomiphene detection task. In all conditions, we observe excellent results, with accuracies over 90% in almost every case. However, we know that the solution to this problem is the appearance of a single additional molecule and its fragments in the spectra, since a solution of water and clomiphene is added in the plasma samples. Thus, it is expected that a single peak (feature) should be sufficient to classify the spectra. Considering this information, we see that a single peak is used for classification only when applying the VLM correction and alignment algorithms when using the Decision Tree and SCM. We also see a decrease in the number of features used for the AdaBoost classifier when using the VLM correction and alignment algorithms. In the case of the L1-regularized SVM, the sparsest solution (with an average of 2.6 features used) was obtained when the VLM correction algorithm was applied in addition to binning.

**Table 1.** Machine learning results in the transductive setting. The percentage in each column is the average accuracy of classifiers on 10 repeats of the experiment. The number shown in parentheses is the average number of features used by the classifiers. The algorithms tested were AdaBoost, the Decision Tree algorithm, the Set Covering Machine (SCM) and a L1-norm Support Vector Machine (L1-SVM).

| Condition | AdaBoost | Decision Tree | SCM | L1-SVM |
| --- | --- | --- | --- | --- |
| Clomiphene Detection |  |  |  |  |
| Binning only | 98.0% (4.7) | 98.6% (1.8) | 95.2% (1.1) | 89.6% (52.0) |
| VLM + Binning | 98.2% (4.9) | 97.0% (2.3) | 97.0% (1.2) | **93.6% (2.6)** |
| VLM + Alignment | **98.8% (2.3)** | **99.4% (1.0)** | **99.4% (1.0)** | 92.8% (138.6) |
| Acetaminophen Detection |  |  |  |  |
| Binning only | 99.2% (1.0) | 99.2% (**1.0**) | 99.2% (1.2) | 97.6% (97.5) |
| VLM + Binning | 99.2% (**1.0**) | 99.2% (**1.0**) | **99.4% (1.0)** | 99.0% (121.0) |
| VLM + Alignment | **99.8% (1.0)** | **100.0% (1.0)** | **99.4% (1.0)** | **99.6% (63.4)** |
| Malaria Detection |  |  |  |  |
| Binning only | 92.4% (51.8) | 82.5% (**4.3**) | 84.6% (2.2) | 92.6% (150.1) |
| VLM + Binning | 93.3% (**39.7**) | **88.7%** (4.6) | **89.4% (2.0)** | **95.4%** (133.2) |
| VLM + Alignment | **93.8%** (65.3) | 86.1% (4.8) | 85.4% (2.3) | 95.2% (**131.4**) |
| Cancer Detection |  |  |  |  |
| Binning Only | **70.4%** (69.2) | **63.8%** (6.4) | 55.6% (**1.9**) | 56.8% (**113.6**) |
| VLM + Binning | 70.2% (43.9) | 61.6% (4.8) | 53.6% (2.2) | 69.4% (138.6) |
| VLM + Alignment | 67.4% (**30.0**) | 62.6% (**2.3**) | **59.6%** (2.2) | **74.6%** (135.2) |

Consider now the results for acetaminophen detection. In this case, an acetaminophen pill was added to the blood plasma samples. Thus, it is expected that multiple molecules and their fragments appear in the spectra in this case, at extremely high concentration not normally found in physiological blood plasma. It is then not surprising that most algorithms can identify acetaminophen with the use of a singe feature (peak). Note that in the case of the L1-regularized Support Vector Machine, the best results, both in terms of accuracy and sparsity, are obtained when the VLM correction and alignment algorithms were used.

The next two tasks represent more realistic problems with unknown solutions. Let us then consider the malaria detection task. For each algorithm, applying the VLM correction algorithm yields an increase in prediction accuracy. For the AdaBoost classifier, we observe an increase of about 1% and the best sparsity in the case of the VLM correction applied before binning, with a slight increase in accuracy with the alignment algorithm. The Decision Tree classifier increases its accuracy by approximately 5% with the VLM correction algorithm, both with alignment and with binning. We see a similar increase in accuracy for the Set Covering Machine in the case of VLM correction with binning. Finally, the L1-regularized SVM obtains a 3% increase in accuracy with the VLM correction algorithm applied, and a better sparsity.

Finally, let us consider the results for the cancer detection task. This classification problem is much harder, with few machine learning algorithms having a prediction accuracy over 70%. Still, both the AdaBoost and Decision Tree classifiers have similar results in all cases, with slight losses in accuracy but improved sparsity with the proposed algorithms applied. The Set Covering Machine sees its accuracy increased by 4% with both correction and alignment algorithms applied and with comparable sparsity. However, in the case of the L1-regularized SVM, the classifier accuracy increases of almost 20% with the proposed algorithms compared to binning only.

### Results for inductive learning

In Table 2, we compare the effect of using the proposed algorithms in the transductive setting versus the inductive setting. For the compound detection tasks, there is very little difference between the two approaches for both clomiphene detection and acetaminophen detection. The inductive setting yields slightly sparser classifiers, but the results are very similar. For the malaria detection task, the difference in sparsity is not significant for the Decision Tree and Set Covering Machine algorithms. The AdaBoost classifier is sparser for the inductive setting, while the L1 SVM has a significant advantage in the transductive setting. The results are also very similar in terms of accuracy for both settings, with very slightly better accuracies in the transductive setting. Finally, the transductive setting appears to be the best setting for cancer detection. The AdaBoost classifier is sparser in this case, with a slight decrease in accuracy. The Decision Tree and Set Covering Machine have better accuracies in the transductive setting, though the SCM is sparser in the inductive setting. The L1-regularized SVM is, on the other hand, much more accurate and slightly sparser in the transductive setting, with an increase in accuracy of about 6%.

**Table 2.**
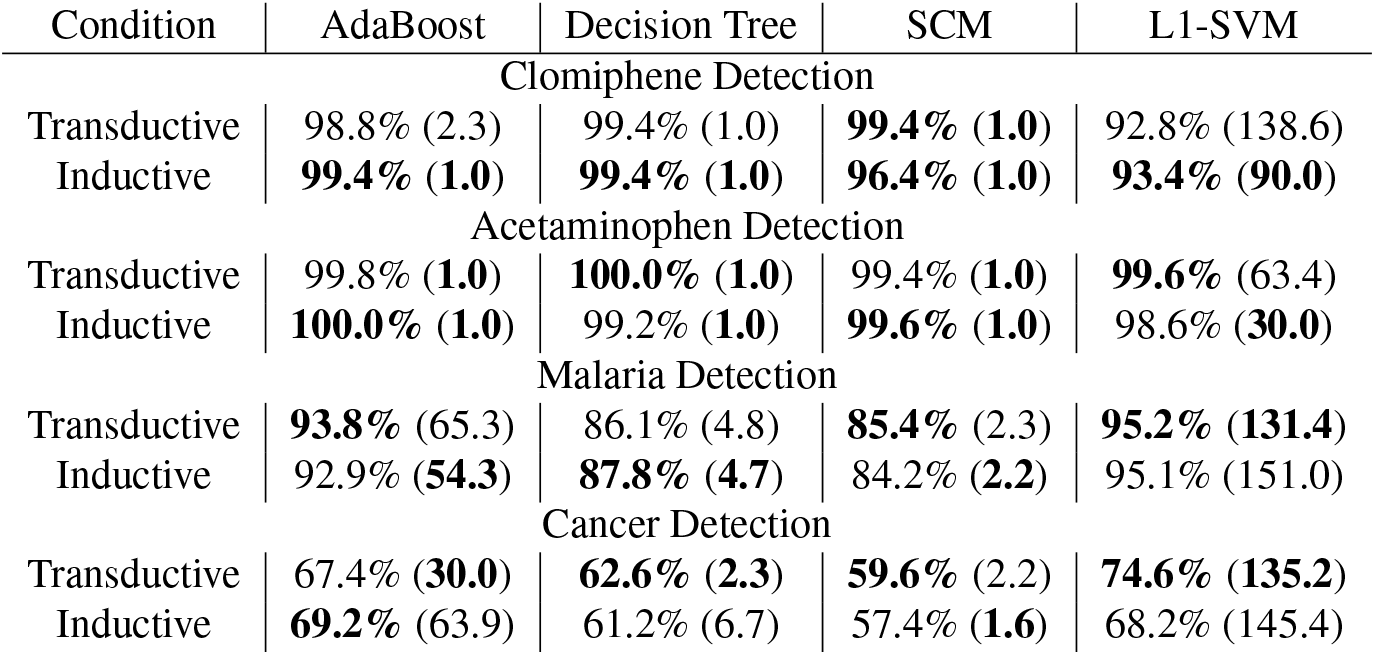
Comparison of transductive and inductive learning of the VLM and Alignment algorithms. The algorithms tested were AdaBoost, the Decision Tree algorithm, the Set Covering Machine (SCM) and a L1-norm Support Vector Machine (L1-SVM).

Finally, and perhaps not surprisingly, we can see (for AdaBoost and L1-SVM) that cancer and malaria detection need far more features then clomiphene and acetaminophen detection.

### Stability of virtual lock masses in datasets

This experiment was conducted in order to verify that virtual lock masses detected on a given dataset will be found in unseen spectra of the same type. The algorithm for VLM detection was also cross-validated on the *Days* Dataset and the *Clomiphene-Acetaminophen* Dataset. Each dataset was randomly partitioned into *k* folds. The VLM detection algorithm was applied to the first *k* − 1 folds, the training folds. The detected VLMs on the training folds are then used for VLM correction of the spectra in last remaining fold, the testing fold. When the correction is applied, we note if every VLM is found in the spectra of the testing fold. The algorithm is scored according to the ratio of detected VLMs on the training folds that are also found in the testing fold. This process is repeated *k* times so that each fold serves as a test fold once. Multiples values of *k* were used in the experiment, such that *k* ∈ {3, 5, 8, 10, 15, 20}.

In each case, we found that *every* VLM point detected on the training set was detected on the testing set. This thus results in a ratio of VLMs found in the testing set over the VLMs detected on the training set of 100% in all cases. This provides empirical evidence of the stability of VLM points across different sets of spectra.

## Discussion

The algorithms proposed in this article aim to render mass spectra more comparable for large datasets acquired in single or multiple batches. The VLM detection algorithm is stable and detects virtual lock masses reliably in datasets. It also detects peaks that are present in mass spectra of the same type but that are not part of the training set. In addition, applying the proposed pipeline of algorithms (VLM detection + VLM correction + alignment point detection) on sets of mass spectra before statistical and machine learning analyses generally yields classifiers with increased accuracy and sometimes with increased sparsity, leading to interpretable models that could serve for biomarker discovery. The proposed pipeline of algorithms has a very low running time complexity of *O*(*n* log *m*) for a collection of *m* spectra containing a total of *n* peaks which, as argued, cannot be surpassed by algorithms based on clustering (with the current state of knowledge). Open-source implementations of the algorithms in Python are also made publicly available.

However, the algorithms, as presented, have a number of drawbacks. Since the virtual lock masses are assigned the average *m*/*z* value of the peaks associated to it, the correction algorithm does not correct the peaks to the exact *m*/*z* value of the ion. The alignment algorithm has also the same property. However, the virtual lock mass approach is compatible with any external lock masses added to the spectra. Thus, by applying both methods, any shift away from the exact (and known) *m*/*z* value of an external lock mass can be corrected. Some situations are also unsuitable for the proposed algorithms. In order for the VLM detection algorithm to function properly and detect virtual lock masses, the mass spectra forming the dataset must be of the same “nature” so that the algorithm can detect a sufficient number of peaks that are common to all spectra. Additionally, the correction algorithm works best in a situation where there are more peaks than spectra. In the cases where each spectrum contains very few peaks, there is a much lower probability that that algorithm can find peaks present in all spectra of the set.

### Future works

The algorithms, as presented here, can only be applied to mass spectra represented by a list of peaks of the form (*μ*, *ι*) where *μ* is the *m*/*z* value of the peak and *ι* its intensity. Hence, the algorithms are currently not applicable with mass spectra having additional dimensions for the peaks, such as ion mobility. It is also not applicable to mass spectra paired with chromatography. It is thus relevant to investigate if the proposed approach, based on virtual lock masses, can be extended to incorporate these extra dimensions.

## Acknowledgements

We thank Waters Corporation for their support and expertise that helped to the design of the proposed algorithms and for the support using their instruments. We also thank Phytronix Technologies Inc. for their support with their instruments. Computations were made on the supercomputer Colosse from Université Laval, managed by Calcul Québec and Compute Canada. The operation of this supercomputer is funded by the Canada Foundation for Innovation (CFI), the ministère de l’Économie, de la science et de l’innovation du Québec (MESI) and the Fonds de recherche du Québec - Nature et technologies (FRQ-NT). Computations were also made on the supercomputer Graham from Waterloo University, managed by Compute Canada. The authors thank the participants for their generosity and providing samples. Plasma samples and the biobanking infrastructure were supported by grants from the Canadian Breast Cancer Research Alliance and the Fondation du cancer du sein du Québec and the Banque de tissus et données of the Réseau de recherche sur le cancer of the Fond de recherche du Québec – Santé (FRQS), associated with the Canadian Tumor Repository Network (CTRNet). DR lab is funded by a CIHR grant MOP130359. DR is a Fonds de la Recherche du Québec-Santé Junior 2 fellow.

## Author contributions statement

F.B., P-L.P., A.D., M.M. and F.L. conceived the algorithm. F.B., M.M. and F.L. conceived the experiments. F.B., P-L.P., A.D. and M.M. programmed the algorithm. F.B. conducted the experiments and analyzed the results. D.G., D.R., F.D. and C.D. validated the results and reviewed the manuscript. J.C. participated to the initial design and reviewed the manuscript. F.B., P-L.P., A.D., M.M. and F.L. wrote and reviewed the manuscript.

## Additional information

### Data Availability

The datasets generated during and/or analysed during the current study are available from the corresponding author on reasonable request.

### Competing Interests Statement

The authors declare no competing interests.

## Notes

#### Summary of Updates

Figures revisited, added an experiment for reviewers. Clarified multiple points. Added contributin authors.

