## Appendix1 for "Mass spectra alignment using virtual lock-masses"

### ABSTRACT

Mass spectrometry is a valued method to evaluate the metabolomics content of a biological sample. The recent advent of rapid ionization technologies such as Laser Diode Thermal Desorption (LDTD) and Direct Analysis in Real Time (DART) has rendered high-throughput mass spectrometry possible. It is used for large-scale comparative analysis of populations of samples. In practice, many factors resulting from the environment, the protocol, and even the instrument itself, can lead to minor discrepancies between spectra, rendering automated comparative analysis difficult. In this work, a sequence/pipeline of algorithms to correct variations between spectra is proposed. The algorithms correct multiple spectra by identifying peaks that are common to all and, from those, computes a spectrum-specific correction. We show that these algorithms increase comparability within large datasets of spectra, facilitating comparative analysis, such as machine learning.

### Supporting information

#### Full Algorithm Description

This section describes, in full detail, the algorithms for Virtual Lock Mass correction and the associated alignment algorithm. Parts of the following sections can be found in the Methods section of the full article. They are repeated here for clarity with the additional details and subsections.

Given a set of $\mathscr{S} = \{S_1, \ldots, S_m\}$ of $m$ spectra, each peak is identified by a pair $(\sigma, \rho)$ where $\sigma \in \{1, \ldots, m\}$ is the index of its spectrum of origin and $\rho \in \{1, \ldots, n_\sigma\}$ is the index of the peak in spectrum $S_\sigma$ containing $n_\sigma$ peaks. Given that we have a total of $n$ peaks in $\mathscr{S}$, we have that $\sum_{\sigma=1}^{m} n_\sigma = n$. For the description of the algorithm, $\mu(\sigma, \rho)$ denotes the $m/z$ value of peak $(\sigma, \rho)$. Finally, we assume that the peaks in each spectra $S_i$ are listed in increasing order of their $m/z$ values.

The proposed algorithm uses two data structures: a *binary heap* and a so-called *active sequence*. A binary heap is a classical data structure used for priority queues which are useful when one wants to efficiently remove the element of highest priority in a queue. In our case, the (binary) heap will maintain, at any time, the next peak of each spectra to be processed by the algorithm. Hence, given a sequence $\mathscr{S}$ of $m$ spectra, the heap generally contains a set of $m$ peaks, where each peak belongs to a different spectrum of $\mathscr{S}$. When a spectrum has no more peak to be processed, the heap contains one less peak. When $k$ spectra have no more peaks to be processed, the heap contains $m - k$ peaks, i.e., one for each spectrum that contains at least one peak to be processed. The "priority value" for each peak $(\sigma, \rho)$ in the heap is given by its $m/z$ value $\mu(\sigma, \rho)$; a peak with the smaller mass is always on top of the heap. That peak can be removed in $O(\log m)$ time for a heap containing $m$ peaks[1]. Given a sequence $\mathscr{S}$ of mass spectra, and a heap $H$ containing at most $m$ peaks, we will use three methods. The first method, $H.init(\mathscr{S})$, constructs the heap that contains the first peak of each spectrum in $\mathscr{S}$ in $O(m)$ time for $|\mathscr{S}| = m$. The second method, $H.top()$, just reads the $m/z$ value of the peak at the top of the heap in $O(1)$ time. Finally, $H.popAndFeed(\mathscr{S})$ removes (and returns to the caller) the peak $(\sigma, \rho)$ on top of the heap, replaces that top element on the heap by peak $(\sigma, \rho + 1)$, which is the next available peak in spectra $S_\sigma$, and then reconstructs the heap in $O(\log m)$ time. Since the peaks in each spectra $S_i$ are assumed to be listed in increasing order of their $m/z$ values, the sequence of calls to $H.popAndFeed(\mathscr{S})$ presents to the caller the peaks from $\mathscr{S}$ in increasing order of their $m/z$ values.

[1]We refer here to the well-known running times (available from any introductory textbook on data structures and algorithms) for heap construction (in $O(m)$ time), removal of its top element (in $O(\log m)$ time), and insertion of a new element (in $O(\log m)$ time).

The second data structure is, what we call, the *active sequence* $A$. At any time, $A$ contains a sequence of peaks, listed in increasing order of their $m/z$ values, which is currently being considered to become a VLM sequence. That data structure uses a doubly linked list $L$ and a boolean-valued vector $B$ of dimension $m$. The linked list $L$ is actually containing the sequence of peaks to be considered for the next VLM and the vector $B$ is such that, at any time, $B[\sigma] = True$ if and only if a peak from spectrum $S_\sigma$ is present in $L$. The active sequence $A$ also maintains the $m/z$ value $\mu_l$ of the last peak that was removed from $L$, the average $m/z$ value $\mu_A$ of the peaks in $L$, and a copy $w_A$ of the window size $w$ chosen by the user. The methods of the active sequence $A$ use the following methods defined for the doubly linked list $L$.

- $L.front()$, which returns a copy of the peak at the beginning of the list $L$ (the one having the smallest $m/z$ value in $L$).
- $L.back()$, which returns a copy the peak at the end of $L$.
- $L.push\_back((\sigma, \rho))$, which inserts the peak $(\sigma, \rho)$ at the end of $L$.
- $L.size()$, which returns the number of peaks in $L$.
- $L.pop\_front()$, which removes the peak at the beginning of the list $L$.

All these methods run in $O(1)$ time for a doubly linked list[2].

The active sequence $A$ provides four methods to the main algorithm. Given an initialized heap $H$ and a chosen window size $w$, the $A.init(H, w)$ method initializes $L$ to be the empty list (containing zero peaks) and initializes all $m$ entries of the vector $B$ to $False$. It also assigns the value $w$ to $w_A$ and $\mu_l$ to a negative $m/z$ value. All these operations in $A.init(H, w)$ are done in $O(1)$ time.

**A.isValid**$(H)$;
**Input:** The heap $H$ that contains at most $|\mathscr{S}| = m$ peaks.
**Output:** $True$ if and only if the set of peaks in $A$ is a VLM point w.r.t. $(\mathscr{S}, w_A)$.
**if** $L.size() \neq m$ **then return** $False$;
**if** $(H.empty() = False) \wedge (\mu(H.top()) \leq \mu_A(1 + w_A))$ **then return** $False$;
**if** $\mu(L.back()) > \mu_A(1 + w_A)$ **then return** $False$;
**if** $\mu(L.front()) < \mu_A(1 - w_A)$ **then return** $False$;
**if** $\mu_l \geq \mu_A(1 - w_A)$ **then return** $False$;
**return** $True$;

**Algorithm 1:** The $A.isValid$ method for active sequence $A$.

The $A.isValid(H)$ method, described by Algorithm (1), returns $True$ if and only if the peaks in the active sequence $A$ satisfies all the criteria enumerated in the definition of a VLM. A precondition for the validity of this method is that $L$ contains only peaks that belong to distinct spectra of $\mathscr{S}$. This precondition holds initially for an empty list $L$ and will always be enforced each time a new peak gets inserted in $A$, as we will see below. Another precondition for the validity of this method is that $\mu_l$ holds the largest $m/z$ value of the peaks in $\mathscr{S}$ that are not in $L$ while satisfying the constraint of being smaller or equal to the $m/z$ value of $L.front()$. This precondition holds trivially initially for an empty list and, as we will see below, will be maintained each time the content of $L$ is modified. The first check of $A.isValid(H)$ returns $False$ immediately as soon as $L$ does not contain exactly $|\mathscr{S}|$ peaks as this violates property (1) for a VLM. Then, if $H$ is not empty, the $m/z$ value of the peak at the top of $H$ (which is the peak having the $m/z$ value closest to the largest $m/z$ value in $L$ while not being smaller to it) is checked to see if it lies outside the interval $[\mu_A(1-w), \mu_A(1+w)]$ permitted for a valid VLM. If it lies inside, we return $False$ as $A$ violates property (4) for a VLM. Note that this check is not performed if $H$ is empty since property (4) is automatically satisfied if there does not exist a peak in $\mathscr{S} \setminus L$ having an $m/z$ value larger or equal to $\mu(L.back())$. The next two checks just verify that all the $m/z$ values of the peaks in $L$ are inside $[\mu_A(1-w), \mu_A(1+w)]$ and return $False$ immediately if this is not the case. The final check verifies if $\mu_l$ lies outside that interval and returns $False$ if it is not the case. Since every check is done in $O(1)$ time, $A.isValid(H)$ executes in $O(1)$ time.

The $A.advanceLowerBound()$ method, described by Algorithm (2) gets called by the main algorithm only when it decides to remove $L.front()$ from the active sequence $A$ in order to eventually include $H.top()$ in $L$. A precondition for its validity, which will be enforced by the main algorithm, is that $L$ is not empty prior to its execution. Before removing the peak $L.front() = (\sigma, \rho)$ from $L$, it first sets $B[\sigma]$ to $False$ since that peak will no longer be in $L$, and assigns $\mu_l$ to $\mu(\sigma, \rho)$. Since no other method removes peaks from $L$, $\mu_l$ is always contains the $m/z$ value of the last peak that moved out from $L$. Then, after the

[2] The fact that these five methods run in $O(1)$ time is the only reason why we have chosen a doubly linked list for $L$. Other data structures such as a circular buffer of size $m$ have also these properties and are, thus, equally valid choices.

**A**.**advanceLowerBound**();
**Output:** Removes $L.front()$ from $L$ and updates the attributes of $A$.
$(\sigma, \rho) \leftarrow L.front()$;
$B[\sigma] \leftarrow False$;
$\mu_l \leftarrow \mu(\sigma, \rho)$;
$L.pop\_front()$;
**if** $L.size() = 0$ **then**
  $\mu_A \leftarrow 0$
**else**
  $\mu_A \leftarrow ((L.size()+1)\mu_A - \mu(\sigma, \rho))/L.size()$
**end**

**Algorithm 2:** The $A.advanceLowerBound$ method for active sequence $A$.

removal of $L.front()$ by $L.pop\_front()$, if it happens that no more peaks are in $L$ it assigns $\mu_A$ to 0 and returns. Otherwise, it sets $\mu_A$ to the average value of the remaining peaks in $L$. Since all these operations are done in $O(1)$ time, this method runs in $O(1)$ time.

**A**.**insert**$(H, \mathscr{S})$;
**Input:** The set of spectra $\mathscr{S}$ and a heap $H$ that contains at most $|\mathscr{S}| = m$ peaks.
**Output:** $True$ if and only if it succeeds at inserting $H.top()$ in the active sequence $A$.
**if** $L.empty()$ **then**
  $(\sigma, \rho) \leftarrow H.popAndFeed()$;
  $B[\sigma] \leftarrow True$;
  $L.push\_back((\sigma, \rho))$;
  $\mu_A \leftarrow \mu(\sigma, \rho)$;
  **return** $True$
**else**
  $(\sigma, \rho) \leftarrow H.top()$;
  **if** $B[\sigma] = True$ **then**
    **return** $False$
  **end**
  $\mu'_A \leftarrow L.size() \times \mu_A + \mu(\sigma, \rho)/(1+L.size())$;
  **if** $\mu(\sigma, \rho) \leq \mu'_A(1+w_A)$ **then**
    **if** $L.front() \geq \mu'_A(1-w_A)$ **then**
      $B[\sigma] \leftarrow True$;
      $\mu_A \leftarrow \mu'_A$;
      $L.push\_back(H.popAndFeed())$;
      **return** $True$
    **end**
  **end**
  **return** $False$;
**end**

**Algorithm 3:** The $A.insert$ method for active sequence $A$.

The $A.insert(H, \mathscr{S})$ method, described by Algorithm (3), tries to insert peak $H.top()$ at the end of $L$. The insertion succeeds if and only if the insertion of $H.top()$ into $L$ gives an $L$ such that the $m/z$ values of $L.front()$ and $L.back()$ (and thus for all other peaks in $L$) lie inside $[\mu_A(1-w_A), \mu_A(1+w_A)]$ and that all peaks in $L$ belong to a different spectra of $\mathscr{S}$. If $L$ is empty when the method is called, then for sure we can insert $H.top()$ into $L$ and, after setting $\mu_A$ to $\mu(H.top())$ and the corresponding entry of $B$ to $True$, we are certain that the desired constraints are satisfied. If $L$ is not empty prior to the insertion of $(\sigma, \rho) = H.top()$, then we verify if another peak coming from spectra $S_\sigma$ is already present in $L$ by testing if $B[\sigma] = True$. If this is the case then we do not insert $H.top()$ and return $False$. If $B[\sigma] = False$ then no peak coming from spectra $S_\sigma$ is in $L$, so we then compute the value $\mu'_A$ that $\mu_A$ should have after the insertion of $(\sigma, \rho)$ and if $\mu(L.front())$ and $\mu(\sigma, \rho)$ both lie in $[\mu'_A(1-w_A), \mu'_A(1+w_A)]$ then we perform the insertion and update $H$ with $L.push\_back(H.popAndFeed())$ and then return $True$. Note that, when we try to insert $H - top()$, we do not verify if $\mu_l < \mu_A(1-w_A)$ with the updated $L$ and $\mu_A$. Indeed, the only fact that we are certain about at all times is that $\mu_l \leq \mu(L.front())$. The purpose of the $A.insert(H, \mathscr{S})$ method is to see if we can insert $H.top()$ into $L$

while still having some probability that the content of $L$ will become a valid VLM sequence after zero or more future insertions. If the insertion is not performed, it means that the content of $L$ with $H.top()$ cannot possibly become a valid VLM sequence and, in that case, to find another VLM sequence we need to remove $L.front()$ with the $A.advanceLowerBound()$ method and then try again to insert $H.top()$ into $L$. No that when the insertion is rejected, $A.insert(H,\mathscr{S})$ is performed in $O(1)$ time since the heap $H$ is not modified. If the insertion is successful, the the heap is reconstructed with $H.popAndFeed()$ in $O(\log m)$ time and, in that case, $A.insert(H,\mathscr{S})$ is performed in $O(\log m)$ time.

**virtualLockMassDetection**$(\mathscr{S},w)$;
**Input:** $\mathscr{S} = \{S_1,S_2,...,S_m\}$, a set of mass spectra.
**Input:** $w$, a window size parameter in relative units.
**Output:** The sequence of all isolated VLM points with respect to $(\mathscr{S},w)$.
**Data:** $H$, a heap initialized with $H.init(\mathscr{S})$; thus containing the first peak of each spectra in $\mathscr{S}$.
**Data:** $A$, an active sequence initialized with $A.init(H,w)$; hence initially empty.
**Data:** $\mathscr{U}$, a sequence of $m/z$ values, initially empty.
$found \leftarrow False$;
**while** $H.empty() = False$ **do**
    **if** $A.isValid(H) = True$ **then** $found = True$;
    **if** $A.insert(H,\mathscr{S}) = false$ **then**
        **if** $found = True$ **then**
            $\mathscr{U}.append(A.get\mu_A())$;
            $found \leftarrow False$;
        **end**
        $A.advanceLowerBound()$;
    **else**
        **if** $H.empty() = True$ **then**
            **while** $A.empty() = False$ **do**
                **if** $A.isValid(H) = True$ **then**
                    $\mathscr{U}.append(A.get\mu_A())$;
                    break;
                **end**
                $A.advanceLowerBound()$;
            **end**
        **end**
    **end**
**end**
**return** $deleteOverlaps(\mathscr{U},w)$;

**Algorithm 4:** The Virtual Lock Mass Detection Algorithm.

Having described the data structures used and their methods, we are now in position to present the main algorithm for virtual lock mass detection, which is described by Algorithm (4). Recall that the task of this algorithm is to find all the isolated VLM points w.r.t. $(\mathscr{S},w)$. The strategy to achieve this task is to first find the sequence $\mathscr{U} = \langle \mu_1,\ldots,\mu_{|\mathscr{U}|}\rangle$ of all possible VLM points w.r.t. $(\mathscr{S},w)$, which may contain several pairs of overlapping VLMs. Once $\mathscr{U}$ is found, Algorithm (4) calls the $DeleteOverlaps(\mathscr{U},w)$ function, described by Algorithm (5), which returns to its caller the subset of $\mathscr{U}$ of all isolated VLM points. For this last task, $DeleteOverlaps(\mathscr{U},w)$ just exploits the definition of overlapping VLM points and, consequently, uses a Boolean-valued vector $A$ (with all entries initialized to $False$) and assigns sequentially $A[k]$ and $A[k+1]$ to $True$ whenever $\mu_k$ overlaps with $\mu_{k+1}$ w.r.t. $w$. Then, to finish its task, Algorithm (5) just needs to insert in $\mathscr{V}$ only the $\mu_k$s for which $A[k]$ is $False$ since it is only these VLMs that do not overlap with another VLM in $\mathscr{U}$. Note that Algorithm (5) runs in $O(|\mathscr{U}|)$ time, which is in $O(n)$ time in the worst case, where $n$ is the total number of peaks in $\mathscr{S}$.

Hence, the central part of $virtualLockMassDetection(\mathscr{S},w)$ is to find the sequence $\mathscr{U}$ of all (possibly overlapping) VLM points. The strategy to achieve this task is to use $A.insert(H,\mathscr{S})$ to try to insert in $A$ (consequently in $L$) the next unprocessed peak of $\mathscr{S}$, which is always located on the top of the heap $H$. Initially, the first such peak of $\mathscr{S}$, a peak having the smallest $m/z$ value among those in $\mathscr{S}$, gets eventually inserted in an empty $A$ by $A.insert(H,\mathscr{S})$. Next, after verifying with $A.isValid(H)$ if the content of $A$ satisfies the criteria to be a valid VLM sequence, we try to insert again in $A$ the next available peak. On each insertion failure, we test if, before this insertion, the content of $A$ was a valid VLM sequence. This is done with the usage of the Boolean variable $found$ (which is set to $True$ as soon as the content of $A$ is a valid VLM sequence and which is set to false immediately after the average $m/z$ value $\mu_A$ of $A$'s content is appended to $\mathscr{U}$). Hence, for each considered peak in $L.front()$, we try to insert one more peak in $L$ and test after the insertion if $L$'s content is a valid VLM sequence. If we cannot insert an extra peak in $A$ with the current peak in $L.front()$ this means that there is no possibility of finding one more VLM sequence with the current peak in $L.front()$. In that case we remove that peak from $L$ with $A.advanceLowerBound()$ and, consequently, $L.front()$ now becomes the peak that was next to $L.front()$ in $L$. Hence, with this strategy, the algorithm attempts to find the largest consecutive sub-sequence of peaks from $\mathscr{S}$ that starts with any given peak in $\mathscr{S}$ and that forms a valid VLM sequence[3]. In addition, note that in the **else** branch of Algorithm (4), we verify if $H$ becomes empty after a successful insertion. In that case, we need to check if we can find a valid VLM sequence by incrementing sequentially the lower bound $L.front()$ and then append to $\mathscr{U}$ the first VLM found. Then, we can safely exit the **while** loop since any other possible VLM sequence will be a subset of the one already found. Without this **else** branch, a VLM sequence that ends with the last peak presented by $H$ would be missed by the algorithm.

For the running time of $virtualLockMassDetection(\mathscr{S}, w)$, note that, at each iteration of the outer **while** loop, the slowest operation is $A.insert(H, \mathscr{S})$, when $H$ is not empty. If $H$ is empty, the slowest operation is the **while** loop in the **else** branch. Since, that operation runs in $O(m)$ time in the worst case and is executed only once, the running time of Algorithm (4) is given by the number of iterations of the outer **while** loop times the running time of $A.insert(H, \mathscr{S})$. Since, for each iteration of the outer **while** loop, either a peak gets inserted into $A$ or removed from $A$, the number of such iterations is in $O(n)$, where $n$ denotes the total number of peaks in $\mathscr{S}$. However, a removal gets performed in $O(1)$ time, whereas an insertion is performed in $O(\log m)$ time, for $m$ spectra in $\mathscr{S}$. Consequently, the running time of Algorithm (4) is in $O(n \log m)$.

```
deleteOverlaps(𝒰, w);
Input: 𝒰 = ⟨μ_1, ..., μ_|𝒰|⟩, a sequence of m/z values (in increasing order).
Input: w, a window size parameter in relative units
Output: The subsequence 𝒱 of m/z values of 𝒰 that do not overlap w.r.t. w.
Data: A, a vector of |𝒰| Booleans initialized to False
Data: 𝒱, a sequence of m/z values, initially empty
for k = 1, ..., |𝒰| − 1 do
    if μ_k(1 + w) ≤ μ_{k+1}(1 − w) then
        A[k + 1] = A[k] = True;
    end
end
for k = 1, ..., |𝒰| do
    if A[k] = False then
        𝒱.append(μ_k);
    end
end
return 𝒱;
```

**Algorithm 5:** Algorithm $deleteOverlaps$

#### An algorithm for Virtual Lock Mass Correction

Given a set $\mathscr{S}$ of spectra and a widow size parameter $w$ expressed in relative units, once the sequence $\mathscr{V}$ of all isolated VLM points w.r.t. $(\mathscr{S}, w)$ has been determined, the individual spectra in $\mathscr{S}$ can be corrected in a manner similar as is usually done with traditional external lock masses. Algorithm (6) performs the correction needed for each peak in a spectrum $S \in \mathscr{S}$. Recall that all the peaks with an intensity smaller than a chosen threshold $t_a$ and greater than a chosen threshold $t_b$ have been removed from $\mathscr{S}$ and that the $m/z$ values in $S$ are listed in increasing order.

First, in the **for** loop, we identify each peak of $S$ corresponding to a lock mass point $v_i \in \mathscr{V}$. Since $S \in \mathscr{S}$ and $v_i$ is a VLM point w.r.t. $(\mathscr{S}, w)$, we are assured to find exactly one such peak $p_j \in S$ with an observed $m/z$ value of $\mu_j$ such that $\mu_j$ lies in the interval $[(1-w)v_i, (1+w)v_i]$. For such $\mu_j$, we assign the index $j$ to $\alpha_i$ so that $\boldsymbol{\alpha} = (\alpha_1, \ldots, \alpha_n)$ is a vector of $n$ indexes, each pointing to the peak in $S$ associated to a VLM point. Note that for $\mu_j \in [(1-w)v_i, (1+w)v_i]$, its corrected $m/z$ value must be equal to $v_i$. Instead of performing these corrections immediately in the **for** loop, we delay them to the linear interpolation step where all peaks having a $m/z$ value $\mu_j$ such that $\alpha_1 \leq j \leq \alpha_n$ will be corrected.

[3] This may appear to be a strategy a bit too complicated than necessary in view of the fact that the largest (and smallest) such sub-sequence must contain exactly $m$ peaks to be a valid VLM. However, we will see below that a significant advantage of using the proposed strategy is the fact that the same algorithm, modulo some very small and trivial modification, can also be used to detect the alignment points of $\mathscr{S}$.

**virtualLockMassCorrection**$(S, \mathscr{V}, w)$;

**Input:** $S = \langle(\mu_1, \iota_1), (\mu_2, \iota_2), ..., (\mu_m, \iota_m)\rangle$, a spectrum

**Input:** $\mathscr{V} = \langle v_1, v_2, ..., v_n\rangle$, a sequence of $m/z$ values (VLMs) sorted in increasing order

**Input:** $w$, a window size parameter in relative units

**Output:** A spectrum $S' = \langle(\mu'_1, \iota_1), (\mu'_2, \iota_2), ..., (\mu'_m, \iota_m)\rangle$ where each $\mu'_j$ is the corrected $m/z$ value for the peak $(\mu_j, \iota_j) \in S$

**Data:** $\boldsymbol{\alpha} = (\alpha_1, \ldots, \alpha_n)$, a vector of indexes (natural numbers)

//construction of $\boldsymbol{\alpha}$

$i \leftarrow 1$;

**for** $j = 1$ **to** $m$ **do**

  **if** $\mu_j \in [v_i(1-w), v_i(1+w)]$ **then**

    $\alpha_i \leftarrow j$ //peak $(\mu_j, \iota_j)$ is associated to VLM $v_i$;

    $i \leftarrow i + 1$;

  **end**

**end**

//correct each $\mu_j$ such that $\alpha_1 \leq j \leq \alpha_n$

$j \leftarrow \alpha_1$;

$i \leftarrow 1$;

**while** $i < n$ **do**

  //linear interpolation correction of $\mu_j$ when $\alpha_i \leq j \leq \alpha_{i+1}$

  $slope \leftarrow \frac{v_{i+1} - v_i}{\mu_{\alpha_{i+1}} - \mu_{\alpha_i}}$ ;

  $b \leftarrow v_i - slope \times \mu_{\alpha_i}$ ;

  **while** $j \geq \alpha_i \wedge j \leq \alpha_{i+1}$ **do**

    //correction of $\mu_j$

    $\mu'_j \leftarrow slope \times \mu_j + b$ ;

    $j \leftarrow j + 1$;

  **end**

  $i \leftarrow i + 1$;

**end**

**Algorithm 6:** Virtual Lock Mass Correction Algorithm

Next, for each VLM $v_i$, we correct by linear interpolation all the $m/z$ values $\mu_j$ such that $\alpha_i \leq j \leq \alpha_{i+1}$. To explain precisely this procedure, let $\mu'(\mu_j)$ denote the corrected value of $\mu_j$. Linear interpolation consists at looking for a correction of the form

$$\mu'(\mu_j) = a\mu_j + b\,,$$

where $a$ is called the *slope* and $b$ is the *intercept*. By imposing that $\mu'(\mu_j) = v_i$ for $j = \alpha_i$ and $\mu'(\mu_j) = v_{i+1}$ for $j = \alpha_{i+1}$, we find that

$$a = \frac{v_{i+1} - v_i}{\mu_{\alpha_{i+1}} - \mu_{\alpha_i}}\,,$$

and $b = v_i - a\mu_{\alpha_i}$. The nested **while** loops of the algorithm performs exactly these linear interpolation corrections for all $\mu_j$ such that $\alpha_i \leq j \leq \alpha_{i+1}$ for $i = 1$ to $n-1$.

Once all $m/z$ values $\mu_j$ such that $\alpha_1 \leq j \leq \alpha_n$ have been corrected, the algorithm is done. Hence, we have decided not to correct any $m/z$ value of $S$ that is either smaller that $v_1(1-w)$ or larger than $v_n(1+w)$ because such a peak has only one adjacent VLM and, consequently, could only be corrected by extrapolation, which is much less reliable than interpolation[4]. Finally, the intensities of the peaks remain unchanged.

For the running time complexity of Algorithm (6), first note that the **for** loop is executed in $O(m)$ time, where here, $m$ denotes the number of peaks in $S$. This is because we only need to iterate once over the peaks in $S$ and, for each peak in $S$, we only perform a constant (independent of $m$ and $n$) amount of operations. Next, the nested **while** loops is also performed in $O(m)$ time since the iteration over all peaks in $S$ is also done only once, and only a constant amount of operations is performed for each peak in $S$. The total running time of the algorithm is therefore in $O(m)$.

#### From VLM correction to spectra alignment

After running the VLM detection and correction algorithms, all the peaks associated with VLM points will be perfectly aligned in the sense that each peak in different spectra associated to a VLM point $v$ will have exactly the same $m/z$ value $v$. However, all the other peaks corrected by Algorithm (6) will not be perfectly aligned in the sense that a molecule fragment responsible for a peak in different spectra will not yield exactly the same mass after correction. This is due to possibly many uncontrollable phenomena that vary each time a sample gets processed by a mass spectrometer, and by the fact that the correction of each peak was performed by an approximate numerical interpolation. However, if all the peaks have been corrected by Algorithm (6), we expect that the peaks corresponding to the same molecule fragment $f$ across different spectra will have very similar masses and will all be localized within a very small window of $m/z$ values. Moreover, we also expect that the $m/z$ values of the peaks coming from another molecule fragment $g$ having a different mass will not cross the $m/z$ values coming from molecule fragment $f$.

More precisely, suppose that we have executed Algorithms (4) and (6) with a window size parameter $w$ (in relative units) on a sequence $\mathscr{S}$ of mass spectra. In addition, suppose that a molecule fragment $f$ gives rise to a peak of $m/z$ value $\mu_1$ in spectrum $S_1$, and a peak of $m/z$ value $\mu_2$ in spectrum $S_2$, and so on for a sub-sequence of spectra in $\mathscr{S}$. Let $\mathscr{M}_f = \langle \mu_1, \mu_2, \ldots \rangle$ be the sequence of these $m/z$ values. Moreover, let $\mu_f$ be the average of the $m/z$ values in $\mathscr{M}_f$. Then, we expect that there exists a window size $\theta$ in relative units, such that $0 < \theta \ll w$, and for which we have $\mu_i \in [\mu_f(1-\theta), \mu_f(1+\theta)]$ for all $\mu_i \in \mathscr{M}_f$. Moreover, if $\theta$ is sufficiently small, we expect that the sequence $\mathscr{M}_g$ referring to peaks produced by another molecule fragment $g$ having a different mass will be such that each $\mu_j \in \mathscr{M}_g$ will not be located within $[\mu_f(1-\theta), \mu_f(1+\theta)]$.

Motivated by this hypothesis, let us introduce the following definitions. Given that Algorithms (4) and (6) have been executed on a sequence $\mathscr{S}$ of mass spectra with window size parameter $w$ in relative units, and given that we have another window size parameter $\theta \ll w$ in relative units, we say that a $m/z$ value $\mu_f$ is an *alignment point* w.r.t. $(\mathscr{S}, \theta)$ if there exists a sequence $\mathscr{M}_f$ of peaks from $\mathscr{S}$ that satisfies the following properties.

1. Every peak in $\mathscr{M}_f$ comes from a different spectrum of $\mathscr{S}$.
2. The average of the $m/z$ values of the peaks in $\mathscr{M}_f$ is equal to $\mu_f$.
3. Every peak in $\mathscr{M}_f$ has an $m/z$ value in $[\mu_f(1-\theta), \mu_f(1+\theta)]$ and all other peaks of $\mathscr{S}$ have an $m/z$ value outside this interval.
4. There does not exist another peak in $\mathscr{S}$ that we can add to $\mathscr{M}_f$ and still satisfy the above properties.

Whenever these criteria are satisfied, we say that $\mathscr{M}_f$ is the *alignment sequence associated* to alignment point $\mu_f$. Given $\mathscr{S}$ and $\theta$, an alignment point $\mu_f$ w.r.t. $(\mathscr{S}, \theta)$ is said to *overlap* with another alignment point $\mu_g$ w.r.t. $(\mathscr{S}, \theta)$ if and only if there exists a nonempty intersection between the $m/z$ intervals $[\mu_f(1-\theta), \mu_f(1+\theta)]$ and $[\mu_g(1-\theta), \mu_g(1+\theta)]$.

[4] Therefore, we recommend removing all these peaks from $S$ to perform statistical analyses or machine learning experiments.

Let $m \stackrel{\text{def}}{=} |\mathscr{S}|$. Notice that the only difference between the definition of alignment point (and its associated alignment sequence) and the definition of VLM point (and its associated VLM sequence) is the fact that a VLM sequence must contain exactly $m$ peaks, whereas an alignment sequence can contain any number of peaks between 1 to $m$ (since the peaks in an alignment sequence may originate from a molecule fragment which is not present in all the samples for which we have a spectrum in $\mathscr{S}$). Hence, if we remove the statement

$$\textbf{if } L.size() \neq m \textbf{ then return } False$$

from the $A.isValid(H)$ method, Algorithm (4) then finds all the maximum-length sub-sequence of peaks that satisfy the 4 criteria for a valid alignment sequence when it reaches the $deleteOverlap(\mathscr{U})$ method. Consequently, with that very minor change,

$$\textbf{virtualLockMassDetection}(\mathscr{S}, \theta)$$

finds all isolated alignment points w.r.t. $(\mathscr{S}, \theta)$ in $O(n \log m)$ time, where $n$ is the total number of peaks in $\mathscr{S}$.

If the window size parameter $\theta$ is too large, then many alignment points will overlap and Algorithm (4) will return very few isolated alignment points. If $\theta$ is very very small, then, in contrast with the VLM identification case, Algorithm (4) will return a very large number of isolated alignment points associated to aligned sequences that contain only one point. Hence, in contrast with the VLM identification case, the best parameter $\theta$ is not the one for which we obtain the largest number of alignment points. What should then be the choice for $\theta$? To answer this question, we consider each VLM point (and its associated sequence of peaks) found by Algorithm (4). If we leave out one VLM point $v_i$ from the correction algorithm (6) and use this algorithm to correct all the $m/z$ values of the peaks associated to this VLM point, the maximum deviation from $v_i$ among these $m/z$ values will give us the smallest window size $\theta_i$ such that each $m/z$ value will be located within $[v_i(1-\theta_i), v_i(1+\theta_i)]$. Essentially, this window size $\theta_i$ is the smallest one for which we can still recognize all the peaks associated to the same VLM $v_i$. It would then certainly be a very good choice for $\theta$ in that region of $m/z$ values. We can then repeat this procedure for all isolated VLM points (except the VLMs with the smallest and largest $m/z$ values) found by Algorithm (4) to obtain a sequence of $\theta_i$ values. One interesting possibility for $\theta$ is the maximum among the $\theta_i$ values. However, this is clearly an overestimate of the maximum spreading of peaks associated to the same molecule fragment since all the VLMs will be used for the correction, including the one that was left out. Moreover, as we can see in Figure (1), we can recover a large fraction of the non-overlapping VLMs if we use a significantly smaller window size than the $\max_i \theta_i$. For that reason, we have decided to use, for the window size $\theta$, the smallest value covering 95% of the non-overlapping VLMs, i.e., the 95th percentile. Alternatively, to attempt to maximize the accuracy of a learning algorithm, a percentile $z$ can be selected by cross-validation along with the selection of the hyperparameters of the learning algorithm.

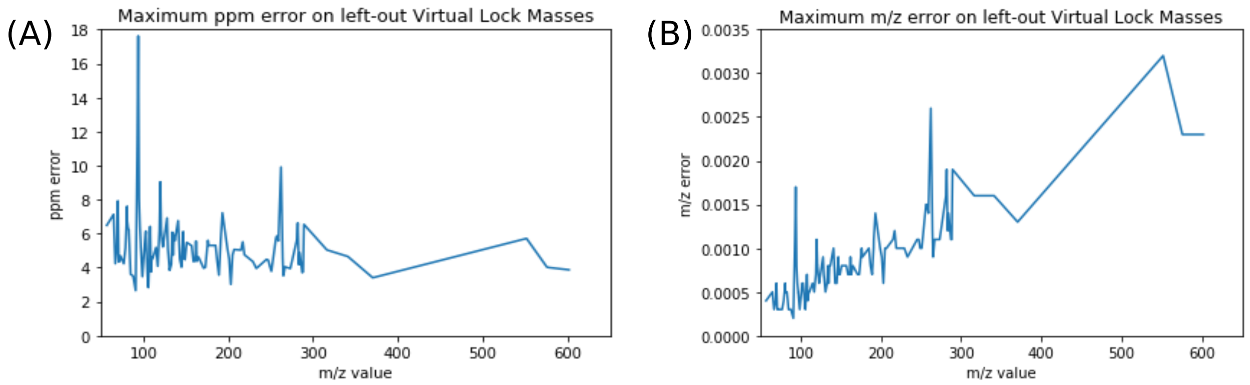


**Figure 1. Error in ppm versus mass units**. Subfigure (A) shows the error on left-out VLMs in ppms, while Subfigure (B) shows the error in Daltons. This data was acquired on the Days Dataset.

If we have $r$ VLM points, each $\theta_i$ associated to the $i$th VLM point is found in $O(m)$ time for a sequence $\mathscr{S}$ of $m$ spectra; thus implying a running time in $O(mr)$ to find every $\theta_i$. Then, the 95th percentile is found by sorting the vector of $\theta_i$s in $O(r \log r)$ time. Assuming that we always have $\log(r) < m$, the total running time to find $\theta$ is in $O(mr)$, and hence in $O(n)$ when $\mathscr{S}$ contains a total of $n$ peaks.

Once the window size $\theta$ is found, we can then run Algorithm (4) just once on the full sequence $\mathscr{S}$ of spectra with that value of $\theta$ in $O(n \log m)$ time. Consequently, the total running time of the alignment algorithm, which includes the running time to find $\theta$ and to find all non overlapping alignment points w.r.t. $(\mathscr{S}, \theta)$, is in $O(n \log m)$.
